# Loss of circulating CD8α^+^ NK cells during human *Mycobacterium tuberculosis* infection

**DOI:** 10.1101/2024.04.16.588542

**Authors:** Nezar Mehanna, Atul Pradhan, Rimanpreet Kaur, Theodota Kontopoulos, Barbara Rosati, David Carlson, Nai-Kong Cheung, Hong Xu, James Bean, Katherine Hsu, Jean-Benoit Le Luduec, Charles Kyriakos Vorkas

## Abstract

Natural Killer (NK) cells can recognize and kill *Mtb*-infected cells in vitro, however their role after natural human exposure has not been well-studied. To identify *Mtb*-responsive NK cell populations, we analyzed the peripheral blood of healthy household contacts of active Tuberculosis (TB) cases and source community donors in an endemic region of Port-au-Prince, Haiti by flow cytometry. We observed higher CD8α expression on NK cells in putative resistors (IGRA-contacts) with a progressive loss of these circulating cells during household-associated latent infection and disease. In vitro assays and CITE-seq analysis of CD8α^+^ NK cells demonstrated enhanced maturity, cytotoxic gene expression, and response to cytokine stimulation relative to CD8α^-^ NK cells. CD8α^+^ NK cells also displayed dynamic surface expression dependent on MHC I in contrast to conventional CD8^+^ T cells. Together, these results support a specialized role for CD8α^+^ NK cell populations during *Mtb* infection correlating with disease resistance.

## INTRODUCTION

Tuberculosis (TB) remains the leading global cause of death from a single bacterial infection and is a significant public health concern (*1, 2*). Exposure to *Mycobacterium tuberculosis* (*Mtb*) bacilli can result in a broad spectrum of clinical outcomes ranging from clearance of primary infection, asymptomatic latent TB infection (LTBI)—measured by *Mtb* peptide-specific interferon γ release assay (IGRA)—or symptomatic active pulmonary TB disease (*2*). It is estimated that 25% of the global population has evidence of latent infection and 10% will progress to active pulmonary disease (*3*). The host immunologic determinants of these clinical outcomes remain poorly defined and are a major obstacle to designing effective TB vaccines and host-directed therapies to prevent disease. Most TB immunology studies focus on conventional peptide-specific T cells (*4–8*), but the presence of these cells have not been shown to correlate with protection (*9–14*). In contrast, few studies have examined *Mtb*-reactive innate lymphocytes that contribute to *Mtb* clearance (*15–18*).

Here, we focus on the most abundant innate lymphocytes in mammals: Natural Killer (NK) cells (*19*). NK cells are prevalent in the peripheral blood (0.1-18% of leukocytes) and populate tissue sites such as liver, lung, and mucosal barriers (*20*). They express a broad range of germ-line encoded pattern recognition receptors enabling rapid responses to infection without the need for antigen-specific expansion (*21*). Prior work provides mechanistic evidence for NK cell recognition of intracellular and extracellular *Mtb* in vitro through natural cytotoxicity receptor engagement of *Mtb* cell wall components or stress-induced self-ligands (*22–33*). However, few published studies directly examine NK cell biology during natural *Mtb* transmission in humans (*27, 28, 34*).

In this study, we interrogate peripheral blood NK cells in a cohort of household TB contacts in an endemic region in Port-au-Prince, Haiti. We analyzed samples from active TB cases (n=50) and their contacts from the same households (n=50) in addition to interferon-gamma (IFNγ) release assay (IGRA)-matched unrelated donors from the source community without household exposure (n=50). We used spectral flow cytometry to test our hypothesis that specific NK cell subpopulations are activated and expand during household-associated *Mtb* exposure and infection. We identify preserved frequency of CD8α^+^ NK cells in blood of uninfected household contacts (IGRA^-^) with progressive loss of this population during household latent infection and active TB disease. CD8α^+^ NK cells demonstrated enhanced responsiveness to cytokine stimulation in vitro compared to CD8α^-^ NK cells and exhibited dynamic surface expression in culture and after in vitro stimulation. Cellular indexing of transcriptomes and epitopes (CITEseq) revealed largely overlapping transcriptional programs between CD8α^+/−^ NK cells with enhanced expression of select cytotoxicity-associated genes in the CD8α^+^ subset. We also observed ubiquitous intracellular NK cell CD8α protein expression. Taken together, our results identify a novel subpopulation of *Mtb*-reactive CD8α^+^ NK cells associated with resistance to primary tuberculosis infection.

## RESULTS

### Clinical cohort description

To study early events during the immune response to *Mtb*, we used biobanked samples from a cross-sectional study performed at GHESKIO Centers, Port-au-Prince, Haiti to assemble a clinical cohort of household contacts of active pulmonary TB cases between 2015-2018 (NIH U19 AI111143, completed). Out of the total 92 household contacts and 591 community controls enrolled, we randomly sampled an age-, sex-, and IGRA-matched subset of 50 TB household contacts and 50 active TB cases from the same households, as well as 50 source community controls without household exposure, based on peripheral blood mononuclear cell (PBMC) availability and quality control (total N=150). All available specimens from IGRA-contacts were analyzed.

Age, sex, IGRA status, and relevant sociodemographic variables are summarized in Table 1. No significant variations were observed in the number of household members, household income, smoking status, or alcohol consumption between groups, consistent with previous work (*15*). Fifty percent of contacts slept in the same room as the active pulmonary TB index case. In the overall study cohort, there were a 50% prevalence of IGRA-positivity in the community controls relative to 75% in contacts, consistent with increased transmission within the household. Among the eleven IGRA-contacts enrolled, only 2 IGRA conversions were detected during 6-month follow-up IGRA testing.

**Table 1.** Demographics of study participants in Haitian Cohort (N=150)

| Variable | Control (n=50) | Contact (n=50) | Active TB (n=50) |
| --- | --- | --- | --- |
| Age – yr | 30.1 ± 9.3 | 33.9 ± 17.8 | 20.2 ± 4.26 |
| ≤ 18 yr | 2 (4) | 12 (24) | 18 (36) |
| >18 and <65 | 48 (96) | 35 (76) | 32 (64) |
| ≥ 65 yr | 0 (0) | 3 | 0 (0) |
| Female sex – no. (%) | 26 (52) | 30 (60) | 26 (52) |
| IGRA positive – no. (%) | 25 (50) | 34 (68) | 38 (76) |

### IGRA+ community controls have elevated frequency of mature CD56^-^ NK cells

We first assessed the frequency of NK cell subpopulations in the clinical cohorts using flow cytometry. Human NK cells were defined by the absence of lineage markers for T cells (CD3), monocytes (CD14), and B cells (CD19), and gated on surface expression of neural cell adhesion molecule (NCAM), CD56, and Fc receptor γRIIIA, CD16 (**Fig. 1A**). We stratified NK cells into three subsets based on expression of CD56: CD56^-^, CD56^dim^, and CD56^bright^ (*35, 36*). There was no difference in frequency of total NK cells or CD56^dim^ NK cells between clinical groups (**Fig. 1B and 1C center**). IGRA+ community controls had fewer CD56^bright^ NK cells (3.11% vs 5.45%, p=0.006) and more CD56^-^ NK cells (19.1% vs 12.6%, p=0.002) compared to IGRA-controls or IGRA+ contacts, indicating an accumulation of mature NK cells during latent TB infection within the source community **(Fig. 1C, right and left)**.

**Fig. 1.**
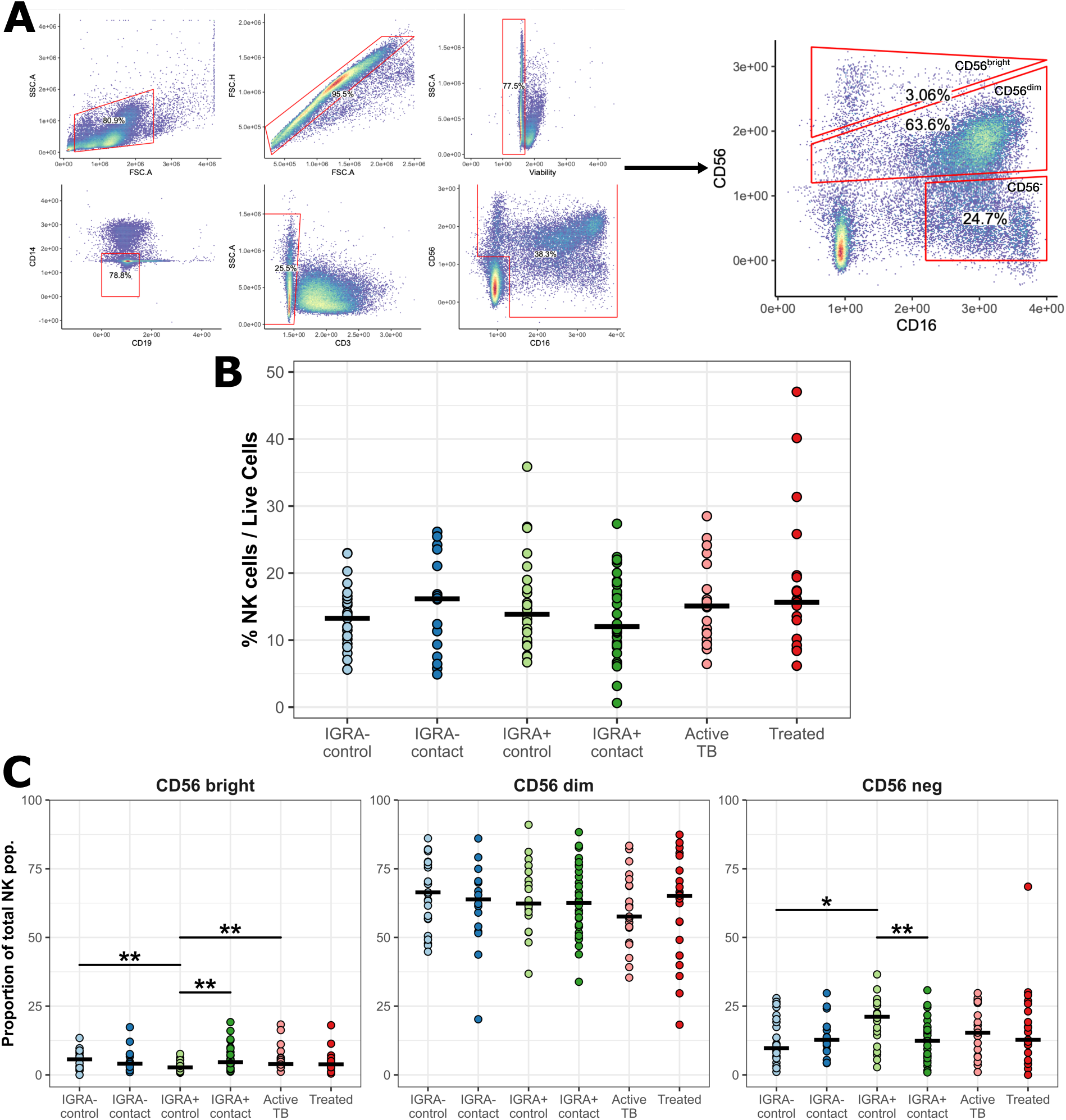
Frequencies of NK cell subpopulations are altered after *Mtb* infection. (**A**) Gating strategy employed for NK cells from a representative sample. NK cells were defined as CD19^-^, CD3^-^, CD14^-^ lymphocytes who expressed any level of either CD16 or CD56. NK cells were further subdivided into three subsets based on CD56 expression. **(B)** Change in frequency of total NK cells as a proportion of total live population. **(C)** Frequency of each NK cell subset in clinical groups. Frequency calculated as percent of each subset among total NK cells. Brackets represent Wilcoxon statistical tests, with unadjusted p-values. *p ≤ 0.05, **p ≤ 0.01.

### IGRA+ contacts demonstrate a loss of circulating CD16^+^ NK cells

NK cell activity against *Mtb* depends upon complex combinatory signaling transduced by activating and inhibiting receptors (*37–39*). We next investigated markers of NK cell activation ex vivo. We analyzed representative receptors, including CD16a (FcγRIIIA), which binds the F_c_ region of antibodies and can induce antibody-dependent cellular cytotoxicity (ADCC). Within the major CD56^dim^ subset, IGRA+ contacts had decreased circulating CD16^+^ NK cells compared to IGRA+ controls (60% vs 80%, p < 0.0001) with a reciprocal increase in CD16^-^ population, representing downregulation or shedding of CD16 surface expression during household-associated LTBI **(Fig. 2A)** (*40*). We found no significant differences in other canonical markers of NK cell activation ex vivo including activation markers CD69 or CD25, or HLA-E restricted receptor NKG2C. We observed decreased staining for degranulation marker CD107a in CD56^bright^ NK cells of IGRA-contacts, potentially representing recent activation events during household exposure. (**Fig. 2B-D**).

**Fig. 2.**
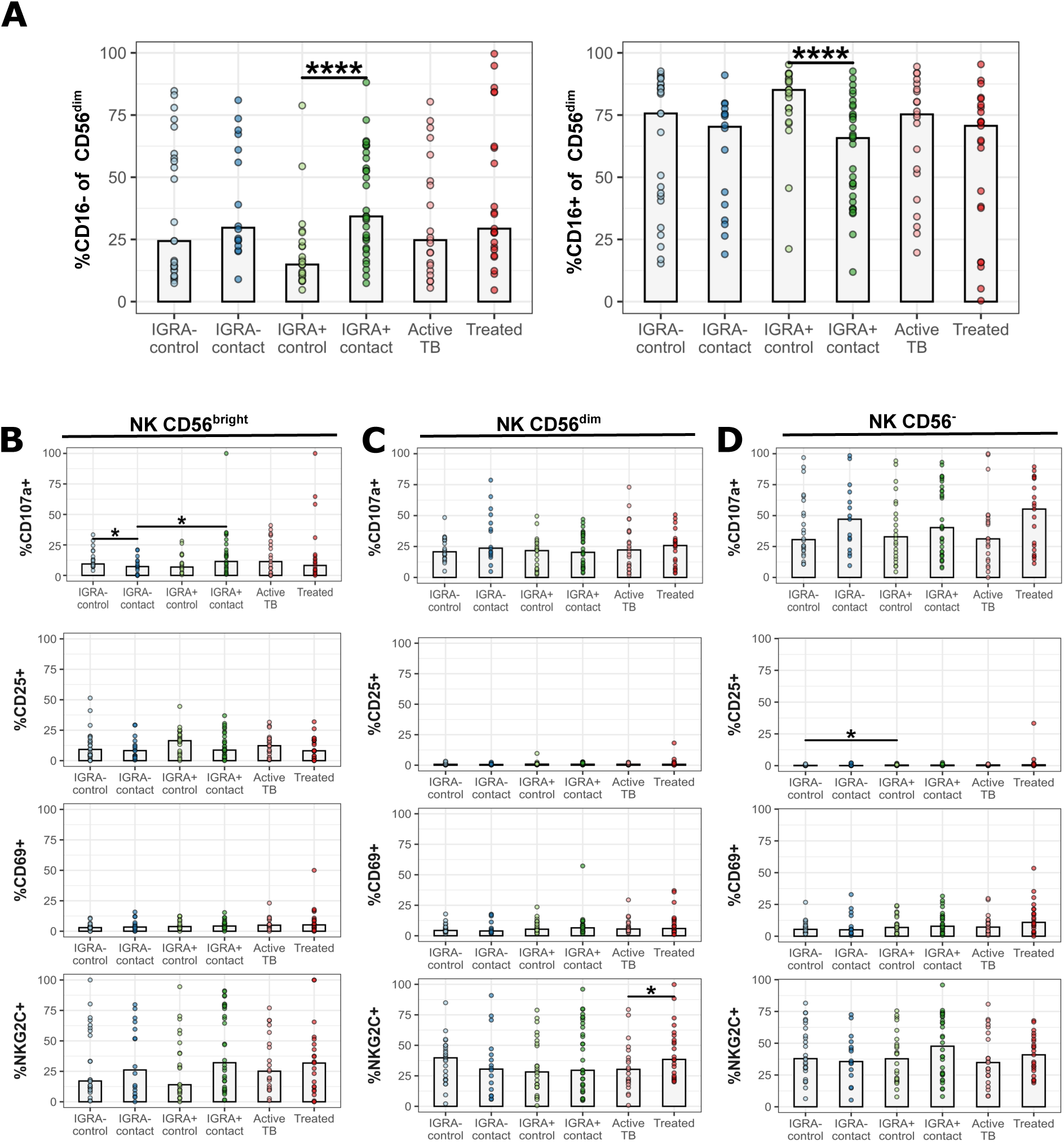
NK cells demonstrate loss of CD16 expression during household-associated latent infection. (**A**) Frequency of CD16 in CD56^dim^ NK cells. Brackets represent Wilcoxon tests with unadjusted p-values. Percent positive for 5 activation markers in **(B)** CD56^bright^ NK cells, **(C)** CD56^dim^ NK cells, and **(D)** CD56^-^ NK cells. Brackets represent Wilcoxon tests with unadjusted p-values. *p ≤ 0.05, ****p ≤ 0.0001.

### NK cells of IGRA-contacts have a depressed functional phenotype

To assess functional activity of NK cells in vitro, we cultured PBMCs in media alone or with K562 cells—a leukemia cell line with low levels of HLA I expression known to activate NK cells—co-incubated at 1:20 K562:PBMC for 15 hours. Cells were then stained, and percent difference was calculated for CD69, CD107a, CD16, and NKG2C. Our positive control analyses in **Fig. 3A** display results from the major CD56^dim^ subset and demonstrate CD107a, CD69, and NKG2C were significantly increased with K562, while CD16 was significantly decreased, across all clinical groups.

**Fig. 3.**
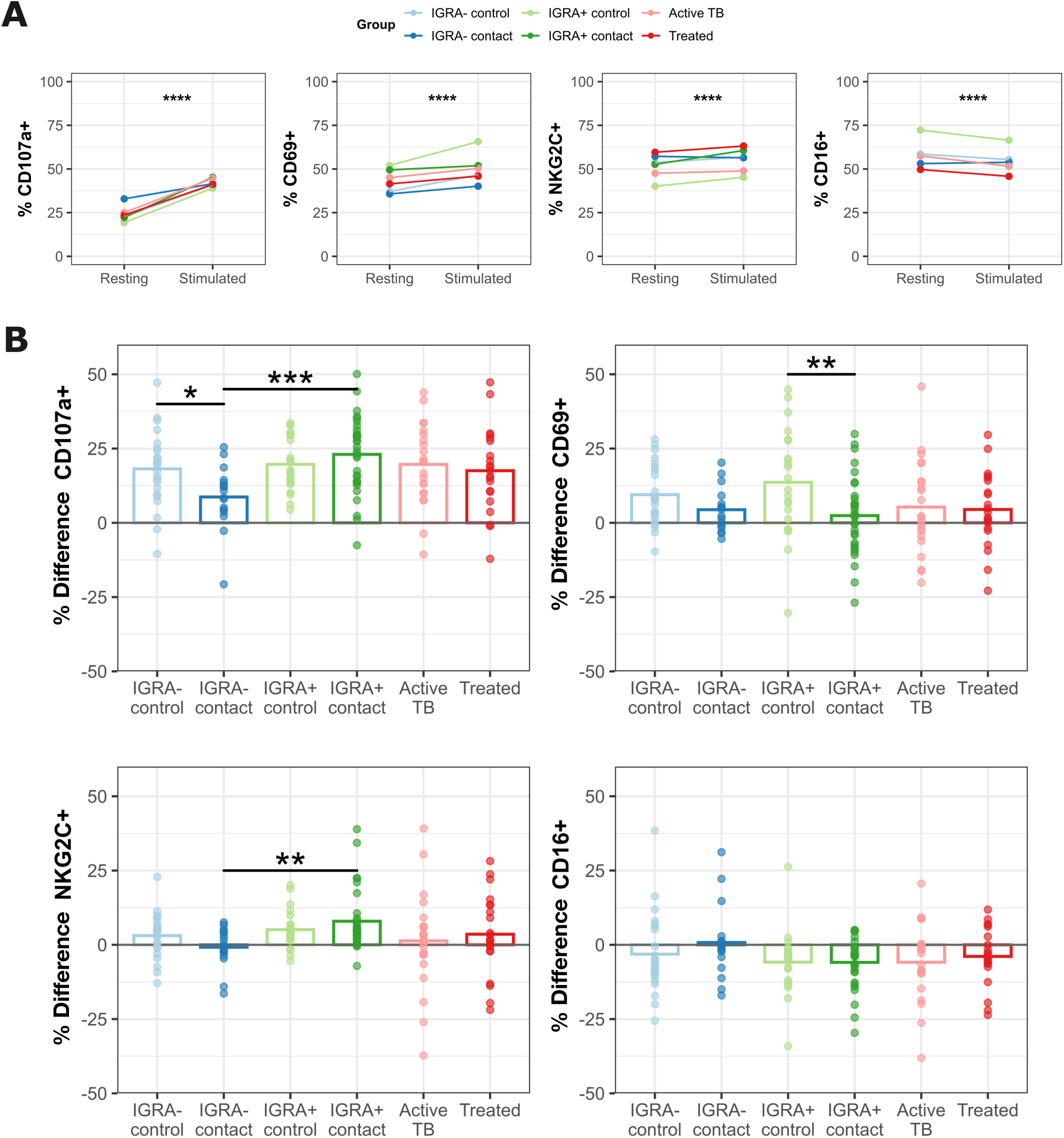
*Mtb* resistors demonstrate depressed NK cell function in vitro. NK cells stimulated overnight by co-culture with K562 cell line compared to plain media alone. **(A)** Overall results of activation assay for CD56^dim^ NK cells for four activation markers. Y-axis represents mean percent positive for each activation marker. P values calculated as a global paired Wilcoxon test including all clinical groups. Each clinical group is displayed separately indicated by line color. **(B)** Percent difference of four activation markers (stimulated minus resting) comparing between clinical groups. Brackets represent Wilcoxon tests with unadjusted p-values. *p ≤ 0.05, **p ≤ 0.01, ***p ≤ 0.001, ****p ≤ 0.0001.

Next, we directly compared NK cell response to K562 between clinical groups. IGRA+ contacts demonstrated significantly more degranulation (CD107a) and NKG2C expression after stimulation relative to IGRA-contacts (**Fig. 3B, left)**. IGRA+ contacts also demonstrated depressed CD69 responses relative to IGRA+ controls (**Fig. 3B, top right**). No significant differences in CD16 expression were detected. The same analyses applied to CD56^-^ and CD56^bright^ NK cells demonstrated similar responses to stimulation, but no significant differences between clinical groups (**Fig. S1, A – D**). Taken together with our ex vivo data, these in vitro restimulation results indicate that CD56^dim^ NK cells of putative *Mtb* resistors (IGRA-contacts) have depressed function, potentially representing sequelae of recent household-associated *Mtb* exposure.

### Loss of circulating CD8α^+^ NK cells during household-associated Mtb infection

Despite their canonical classification as short-lived innate lymphocytes, NK cells are increasingly recognized for their marked diversity and ability to differentiate into longer-lived effectors (*16, 18, 27, 41–44*), including after mycobacterial stimuli (*45*). The most extensively studied is an NK cell memory population co-expressing NKG2C and CD57 that expands in blood during human cytomegalovirus (CMV) infection (*27, 46*). Multiple putative markers of NK cell memory during *Mtb* infection have been suggested, including CD57, CD8α, CD161, and CD27 (*28, 34, 45*). We compared expression of these markers across clinical groups and found that household-associated LTBI (IGRA+ contacts) and active TB disease donors had progressively decreasing frequency of CD8α^+^CD56^dim^ NK cells (**Fig. 4A)**. A similar trend was observed with the marker of terminal differentiation, CD57. There was significantly fewer CD57^+^CD56^dim^ NK cells in household-associated LTBI (IGRA+ contacts) compared to healthy household contacts (**Fig. 4B**). There was no association between CD8α expression and age or sex, and loss of CD8α^+^ was specific to NK cells and was not observed among T cells during active TB (**Fig. S2A-B**). We did not observe any significant differences in the expression of CD27, CD127, or CD161 between clinical groups (**Fig S3**). Together, these results indicate that donors who develop household-associated LTBI and active TB disease experience a loss of circulating CD8α^+^ and CD57^+^ CD56^dim^ NK cells. In turn, CD8α^+^ NK cell frequency significantly correlates with resistance to primary household-associated infection.

**Fig. 4.**
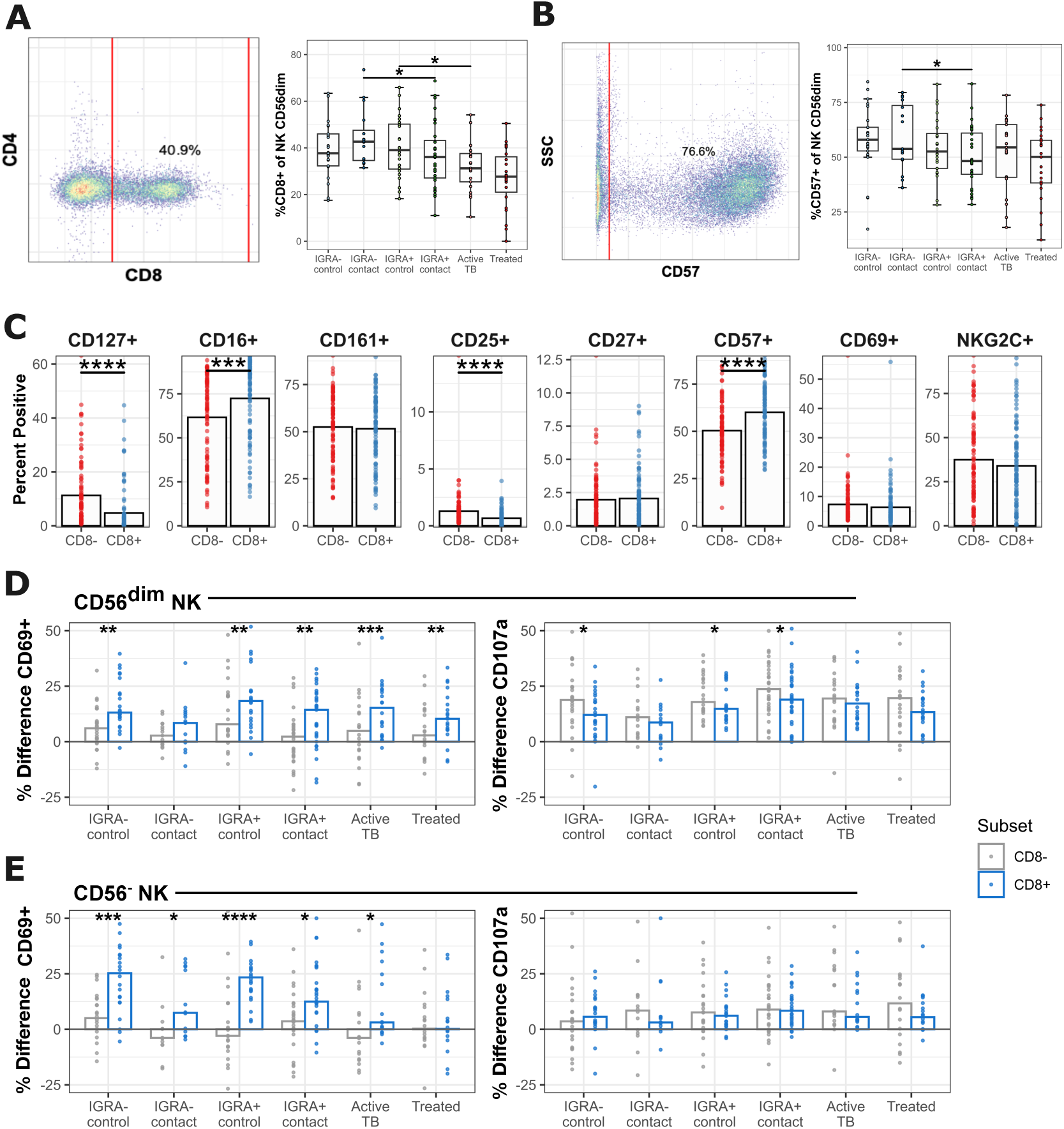
CD8α^+^ NK cells are depleted in the blood of cases with household-associated *Mtb* infection. (**A**) Representative gating of CD8α^+^ in CD56^dim^ NK cells and frequency of CD8α^+^ within CD56^dim^ NK cells. **(B)** Representative gating of CD57 in CD56^dim^ NK cells and boxplots of percent CD57^+^ within CD56^dim^ NK cells. **(C)** Expression of select surface markers between CD8α^+^ vs CD8α^-^ CD56^dim^ NK cells, active TB are excluded. **(D)** Activation assay with overnight K562 co-culture versus plain media comparing CD8α^+^ vs CD8α ^-^ CD56^dim^ NK cells. Y-values represent percent difference for CD69 and CD107a. Asterisks represent paired Wilcoxon tests. **(E)** As in panel D, instead displaying CD56^-^ NK cells. P-values represent unadjusted Wilcoxon testing. *p ≤ 0.05, **p ≤ 0.01, ***p ≤ 0.001, ****p ≤ 0.0001.

### CD8α^+^ NK cells represent a mature population with high functional potential

Despite the identification of cytotoxic CD8α^+^ NK cells over 30 years ago (*47–49*), the function of CD8α on NK cells during health and disease is not well-characterized and is of emerging interest as a cellular target against immune-mediated diseases (*47, 49, 50*). We next used flow cytometry to define co-expressed receptors of CD8α^+^ NK cells in our cohorts. In the Haitian cohorts, a mean of 27% of NK cells expressed CD8α on their surface. Among NK cell subsets, CD56^dim^ NK cells had the largest proportion of CD8 positivity (median: 36%, range: 1-74%) (**Fig. S4A**). We then defined co-expressed receptors on CD8α^+^ NK cells among all healthy Haitian donors, excluding active TB cases with chronic symptomatic disease. CD8α^+^ NK cells expressed significantly higher levels of CD57 and CD16, and lower levels of CD25 and CD127 (**Fig. 4C**). This indicates that CD8α^+^ NK cells represent more mature effectors compared to their CD8α^-^ counterparts. When compared to T cells, median fluorescence intensity (MFI) of CD8α surface expression was significantly lower in NK cells (**Fig. S4B**).

To interrogate function, PBMCs were co-incubated with K562 cells for 15 hours and compared to resting conditions as described above, stratified by CD8α. CD8α^+^ NK cells were more responsive to K562 induction measured by CD69 upregulation compared to CD8α^-^ NK cells but demonstrated lower levels of degranulation as measured by CD107a staining (**Fig. 4D**). The propensity for CD8α^+^ NK cells to upregulate CD69 upon K562 stimulation was more pronounced among the CD56^-^ compartment, where the CD8α^-^ fraction had little to no response (**Fig. 4E**). Overall, this suggests that CD8α expression among NK cells marks a population with higher capacity for activation, specifically among terminally differentiated CD56^-^ NK cells, that are reported to have depressed effector functions (*51*).

### CD8α is ubiquitously expressed in NK cells

We next interrogated the immune phenotype and function of CD8α^+^ NK cells in a distinct biorepository of healthy human PBMCs (IRB2021-00478; PI: Vorkas) to validate our findings and perform additional functional assays. First, we confirmed the presence of CD8α expression in NK cells by extracellular staining with CD8α and CD8β antibodies. About 60% of NK cells did not express surface CD8, CD8αα was found in ∼40% of NK cells, while rare CD8αβ NK cells were detected (**Fig. 5A**). In blood, CD8α was expressed in all three compartments of NK cells, and most frequently among CD56^dim^ cells, with median expression comparable to Haitian donors (**Fig. 5B**). We also validated our prior result from the contact cohort (**Fig. 4C**) demonstrating CD16 and CD8α were significantly co-expressed (**Fig. 5C).** As our TB contact studies only assessed surface CD8α on NK cells, we next analyzed intracellular CD8α expression by flow cytometry. We observed that nearly all CD56^dim^ NK cells expressed CD8α protein intracellularly, contrasting with T cells whose CD8 expression did not significantly differ between surface and intracellular staining (**Fig. 5D**). CD56^bright^ and CD56^-^ NK cells also demonstrated ubiquitous expression of CD8α protein intracellularly (**Fig. S5**). To test the stability of surface CD8α on NK cells over time, we incubated PBMCs for 1 or 7 days in pro-survival cytokine IL-2. We observed significantly increased CD8α surface expression following prolonged incubation, indicating that surface presentation of NK cell CD8α is more dynamic than in T cells where it is a constitutively expressed surface receptor (**Fig. 5E**).

**Fig. 5.**
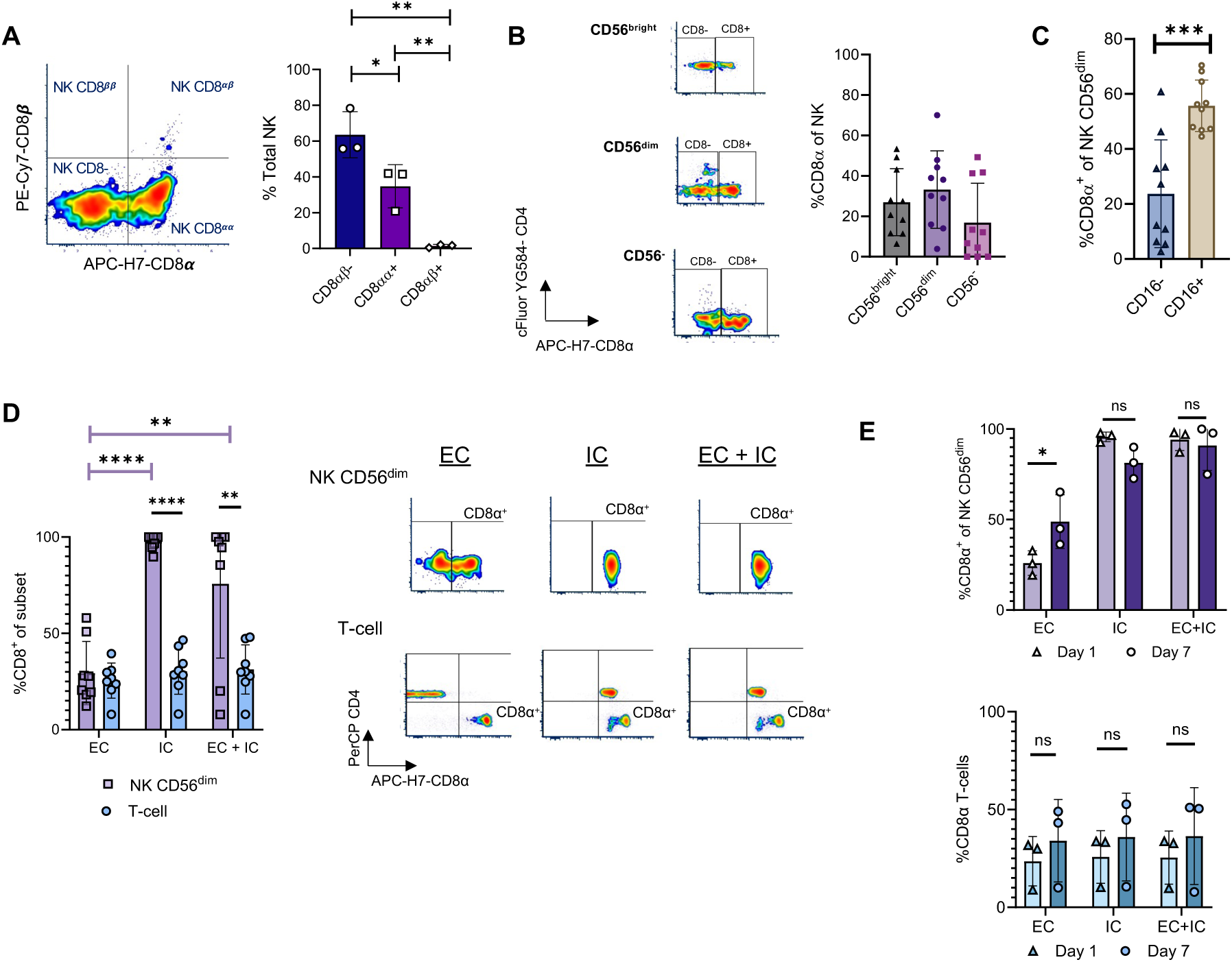
CD8α^+^ NK cells are prevalent in healthy US donors with ubiquitous intracellular CD8α expression. (**A**) Flow cytometry plot and percentage of CD8αα^-^, CD8αα^+^, CD8αβ^+^, and CD8ββ^+^ among total NK (N=3). **(B)** Percentage of CD8α^+^ NK cells in different subsets based on CD56 expression (N=10). **(C)** Frequency of CD16 expression in CD8α^+^ CD56^dim^ NK cells. **(D)** Frequency and representative flow cytometry plots of CD8α expression comparing CD56^dim^ NK cells and T cells with extracellular (EC), IC (Intracellular), and EC + IC CD8α staining. **(E)** Percent CD8α^+^ in EC, IC, and EC + IC staining at day 1 and day 7 in CD56^dim^ NK cells and T cells. Statistical significance was measured using unpaired t tests, ns – no significance, *p<0.05, **p<0.005, ***p<0.001, ****p<0.0001.

### CD8α^+^ NK cells have enhanced responses to cytokine stimulation

We next hypothesized that surface CD8α was associated with enhanced responsiveness to inflammatory signals, following prior reports of increased cytotoxicity against leukemic cells relative to CD8α^-^ NK cells (*48*). We stimulated PBMCs with either the potent IL15-IL15Rα sushi domain fusion construct (IL15-15RαSu) (*52*), IL15-15RαSu/IL12/IL18, IL12/IL18 alone, or virulent *Mtb* Erdman whole cell lysate and assessed intracellular IFNγ expression by flow cytometry. Cytokine stimulation led to decreased CD8α^+^ surface staining while *Mtb* stimulation did not alter CD8 surface expression, suggesting that cytokine stimulation induces downregulation of the CD8α receptor or cell death (**Fig. 6A**). Both IL15/IL12/IL18 and to a lesser degree *Mtb* lysate conditions induced IFNγ responses in CD56^dim^ NK cells compared to resting (**Fig. 6B**). When stratified by CD8α, we observed increased intracellular IFNγ with cytokine stimulation in both CD16^-/+^ CD8α^+^ NK cells relative to CD8α^-^ cells, but no differences upon co-incubation with *Mtb* lysates (**Fig. 6C**). Percentage of CD25, CD69, and intracellular Granzyme B responses were also enhanced in CD8α^+^ NK cells across CD56-stratified subsets (**Fig. S6**).

**Fig. 6.**
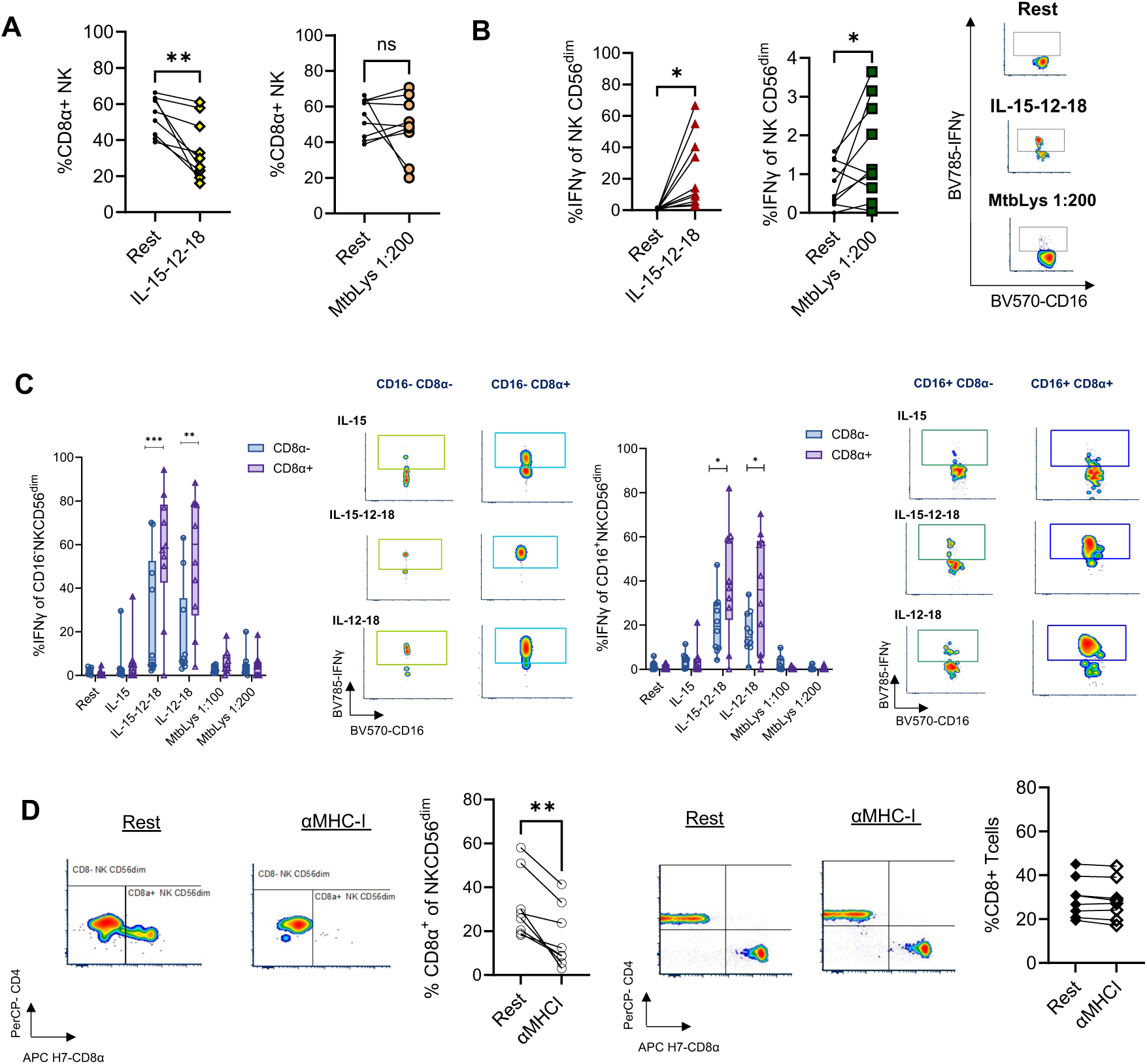
CD8α^+^ CD56^dim^ NK cells have enhanced response to cytokine stimulation. (**A**) Pairwise analysis of percent CD8α^+^ NK cells upon stimulation with IL15/12/18 and MtbLys 1:200 (n=9). (**B**) Pairwise analysis and representative flow plots of intracellular IFNγ in NK CD56^dim^ upon stimulation with IL15/12/18 and MtbLys at dilution 1:200 (n=9). (**C**) Percent intracellular IFNγ^+^ in CD8α^+^ versus CD8α^-^ CD56^dim^ NK cells comparing IL-15-Su, IL15-Su/12/18, IL-12/18, or MtbLys conditions (n=9). Both the CD16^+^ (right) and CD16^-^ (left) CD56^dim^ NK subpopulations are displayed. (**D**) Representative flow cytometry plots and paired analysis of percent change CD8α^+^ with anti-MHC I for 15 hours among CD56^dim^ NK cells (left) and CD8^+^ T cells (right) (n=8). Statistical significance was measured using paired t-test, with threshold of *p<0.05, **p<0.005, ***p<0.001. IFNγ: Interferon γ, Mtb Lys: *Mtb* whole cell lysate.

### CD8α^+^ surface expression on NK cells is MHC I-dependent

To determine a mechanism by which CD8α may modulate NK cell function, we hypothesized that CD8α may engage MHC I molecules like CD8^+^ T cells (*53, 54*). To test this, we stimulated whole PBMCs with MHC I blocking antibody (clone: W6/32) and measured surface CD8α expression. Strikingly, surface CD8α expression on NK cells—but not T cells—was significantly decreased in all anti-MHC I conditions (**Fig. 6D**). Our results indicate that CD8α surface expression is maintained through engagement of MHC I molecules.

### CD8α^+^ NK cells have overlapping transcriptomes with CD8α^-^ NK cells

While CD8α-expressing NK cells have previously been described (*47–49, 55*), the transcriptional programs associated with surface expression of CD8α and whether these relate to homologous function in CD8α^+^ T cells during disease is an area of emerging interest. To interrogate the transcriptional profile of CD8α^+^ NK cells, we sorted innate lymphocyte subsets from PBMCs including NK cells, γδ T cells, MAIT cells, and invariant Natural Killer T cells (iNKTs) by fluorescence-activated cell sorting (FACS) from the peripheral blood of two healthy donors. We performed CITE-seq (*7*) of these populations directly after sorting or after coincubation for seven days in IL2/IL7 supplemented media +/− *Mtb* lysates in order to define surface protein and transcript expression in CD8α NK cells. We discriminated NK cells from surrounding innate lymphocytes based on surface antibody barcodes for CD56 and CD16 together with expression of canonical NK cell gene transcripts *CD3E*, *NCR1*, and *NCAM1* (CD56) (**Fig. S7**) (*47, 56, 57*). Dimensionality reduction with UMAP identified four clusters of NK cells with distinct transcriptional signatures (**Fig. 7, A to D**). To assess pathogen-specific responses, we first performed differential expression between the ‘rest’ and ‘*Mtb* lysate’ conditions and display the top 8 most differentially expression genes in **Fig. 7E**. The cytotoxic serine protease *GZMB (*Granzyme B) and killer immunoglobulin-like receptor, *KIR3DL1*, were the most upregulated genes in the *Mtb* lysate condition.

**Fig. 7.**
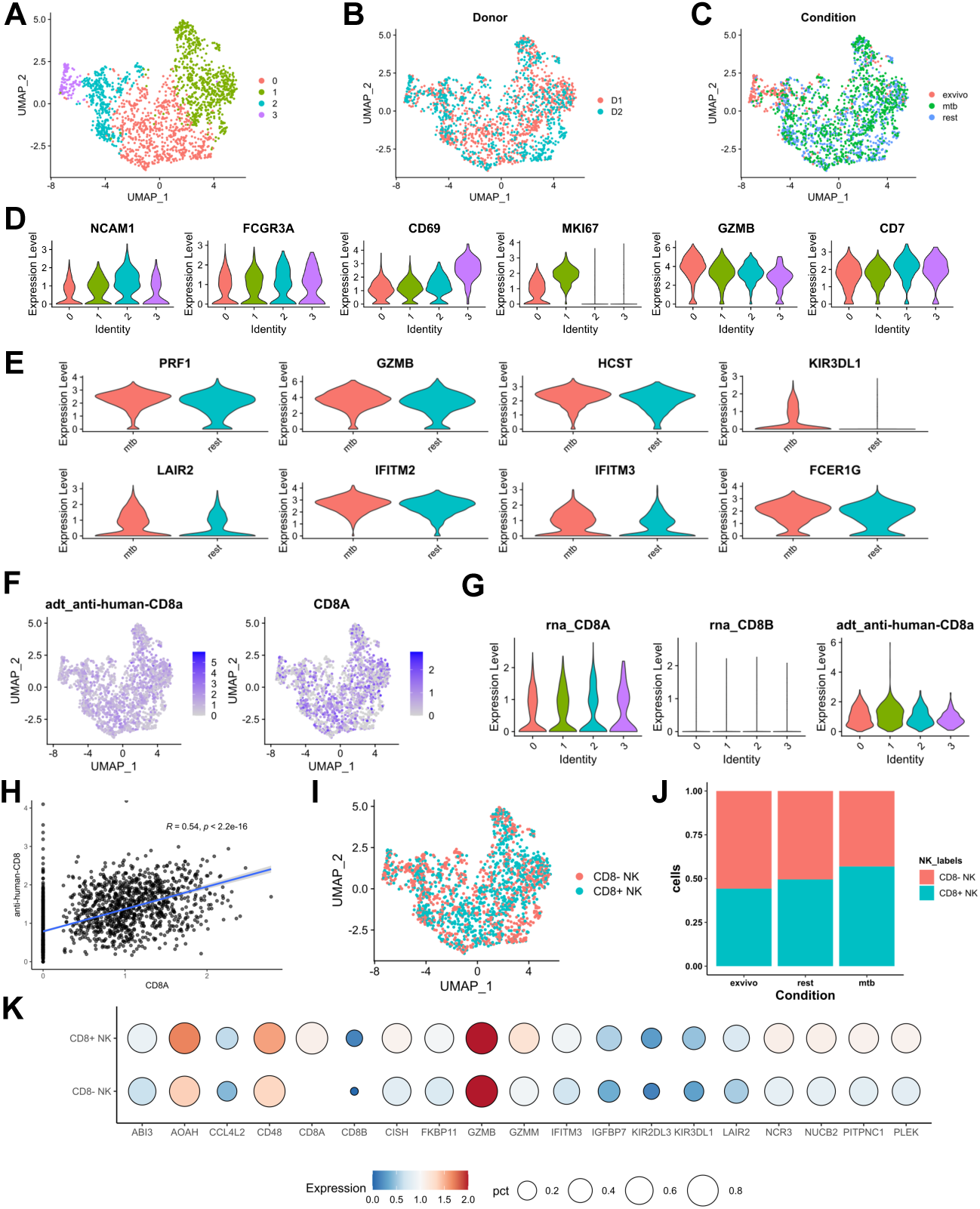
CITE-seq analysis of CD8α^+^ vs CD8α^-^ NK cells from two healthy donors reveals overlapping transcriptomes. (**A**) Results of clustering and uniform manifold approximation (UMAP) projection for all NK cells. **(B)** UMAP reduction colored by Donor. **(C)** UMAP reduction colored by condition. **(D)** Violin plots representing top six genes that were differentially expressed between the four NK cell clusters. **(E)** Violin plots of the eight differentially expressed genes between resting and *Mtb* lysate condition. **(F)** Distribution of NK cell expression of CD8 based on barcoded antibodies against CD8α and *CD8A* transcript. **(G)** Violin plot of the expression of *CD8A* and *CD8B* transcripts, as well as CD8α surface expression between the four NK cell clusters. **(H)** Correlation of CD8α expression between transcript and protein measurement. **(I)** Annotated UMAP based on transcript expression of *CD8A* into CD8α^+^ and CD8α^-^ NK cells. **(J)** Percent of NK cells expressing CD8A transcript between stimulation conditions. **(K)** Dot-plot of genes differentially expressed between CD8α^+^ and CD8α^-^ NK cells.

We next analyzed the distribution of *CD8A* transcript and CD8α protein expression among NK cells using both transcript and surface antibody barcodes for CD8α. NK cells did not cluster independently based on the presence of *CD8A/*CD8α (**Fig. 8, F and G**) and *CD8A*^+^/CD8α^+^ NK cells were distributed throughout all clusters (**Fig. 8I)**. Low amplification of surface antibody barcodes for CD8α was observed in NK cells, in contrast with other CD8α –expressing immune subsets sequenced (e.g., MAIT cells; **Fig. S8**) in line with CD8α MFI differences detected by flow cytometry (**Fig. S4B**).

To define transcriptional signatures associated with *CD8A* transcript expression, we stratified NK cells expressing greater than zero *CD8A* transcripts and performed differential expression relative to *CD8A*^-^ cells. Half of all NK cells expressed *CD8A* transcripts (**Fig. 8J**). Differential gene expression revealed few statistically significant differences between *CD8A^+/^*^-^ cells. Notable differentially expressed genes in *CD8A*^+^ NK cells include cytotoxicity-associated proteins natural cytotoxicity receptor 3 (*NCR3*) and the serine protease granzyme M (*GZMM*) (**Fig. 8K**).

## DISCUSSION

Identifying the correlates of protection against natural *Mtb* infection is an area of critical importance that will inform the next generation of TB vaccines and host-directed therapies (*5, 37, 45, 58–61*). In this study, we hypothesized that investigation of household-associated *Mtb* exposure would reveal specialized NK cell populations and represent early events following high-risk transmission. Our results identify a CD8α-expressing NK cell subpopulation with preserved frequency in the blood of contacts who remain uninfected (IGRA-) that is progressively depleted during household LTBI and active TB disease. Few studies have interrogated CD8α^+^ NK cells, as this receptor is most commonly understood as a lineage marker for T cells (*62*) and is absent in specific pathogen free murine NK cells (*63*). Recent work reporting an association of CD8α^+^ NK cells with protection against human infectious, autoinflammatory and malignant disease has generated interest in targeting this subpopulation for host-directed therapeutics (*39, 47, 48*). However, little is known how CD8α receptors may modulate NK cell activity, namely, enhanced cytotoxic functions previously reported in vitro against leukemia cells and strong associations with protection against both infectious and non-infectious diseases. (*48, 50, 55*). Here, we show that the majority of peripheral blood NK cells can express CD8α protein, but neither *CD8A* transcript nor CD8α protein marked a distinct transcriptional state in healthy donors. Importantly, we demonstrate that surface expression of CD8α on NK cells depends upon MHC I expression and marks a population with enhanced responsiveness to cytokine stimuli in vitro. Together, our study highlights a novel innate effector against human tuberculosis and provides important insights into NK cell biology and innate TB immunology.

To our knowledge, there have been only two published household TB contact studies that specifically examined NK cell biology during early *Mtb* exposure and infection. Our study expands upon the sample size of this prior work and provides additional analyses of NK cell subpopulations stratified by CD56 not previously addressed (*42, 64, 65*). Specifically, we observed decreased frequency of immature CD56^bright^ NK cells with reciprocal increase in mature CD56^-^ NK cells during community LTBI, in line with findings during chronic viral infections such hepatitis C and HIV (*66–68*). We also observed that CD16 expression was increased during community LTBI relative to household-associated infection, which may represent a late effect of latency within the CD56^dim^ subset (*61*). An equally valid interpretation of these results is that loss of surface CD16 is an early event detected after household-associated exposure and may represent downregulation or shedding of the F_c_γ receptor after engagement (*40*). Together, these changes in frequency of CD16 and CD56 expression indicate a gradual expansion of mature NK cells in peripheral blood after asymptomatic infection (*29, 40, 64, 65, 67, 69*), accompanied by both markers of maturity and dysfunction (loss of CD56), but also enhanced potential for ADCC (gain of CD16).

Two phenotypic markers whose surface expression were both independently and inversely associated with infection in households, were CD8α and CD57. CD8α has long been appreciated to be expressed on the surface of human (*47–50, 70*) and non-human primate NK cells (*63, 71*). CD8α expression on human NK cells has been associated with enhanced cytotoxicity (*48, 55*) and is referred to in one study as a putative *Mtb* memory marker (*45*). Recent studies in prospective longitudinal human cohorts report that loss of CD8α expression at the transcript (*CD8A*) or protein level correlate with HIV progression (*39, 49*), relapsing multiple sclerosis (*47*), and recurrence of acute myeloid leukemia after stem cell transplantation (*50*), where CD8α^+^ NK cells are hypothesized to exercise combined cytotoxic and regulatory roles to prevent disease. Sorted surface CD8α^+^ NK cells also demonstrate enhanced anti-leukemia cytotoxicity in vitro compared to sorted CD8α^-^ cells (*50*). We also report that CD8α^+^ NK cells were prevalent in healthy US donors with ubiquitous intracellular CD8α protein expression. This finding suggests that most NK cells are capable of presenting CD8α to the cell surface to enhance effector function, in contrast to constitutive surface CD8α expression on T cells. Moreover, CD8α^+^ NK cells were exquisitely sensitive to cytokine stimulation, suggesting complex roles during inflammatory immune responses against disease. Further, we show that CD8α surface expression was strongly dependent on MHC I expression (*50, 54, 72*) raising critical questions for future studies to define the molecular mechanisms of CD8α engagement with MHC I and to determine whether this may be modulated by different peptide-MHC I complexes. Taken together with previously published work (*47–49*), CD8α^+^ NK cell subpopulations emerge as an attractive target in vaccination and host-directed approaches against infectious and noninfectious diseases (*73*).

In addition to these mechanistic insights, we present the first multi-omic analysis of CD8α^+^ NK cells and show that in contrast to T cells, CD8α does not mark a distinct transcriptional state of NK cells in healthy donors assayed, though *CD8A*^+^ NK cells did differentially express cytotoxic genes including *NCR3* and *GZMM*. These results will need to be validated in other healthy donors as well as compared to future CITE-seq experiments in high-risk *Mtb* exposure cohorts or other disease states.

There are certain limitations inherent to our human immunology study design. While household contacts experienced high risk exposures to pre-treatment smear+ active TB cases, this does not exclude the possibility they may have been exposed or infected in the community. We control for this using source community donors to account for background transmission. Similarly, IGRA-community controls may have also been recently exposed and resisted primary infection, though this group represents the most rigorous control for household exposure. Our study was conducted in one TB endemic population in Haiti and will need to be validated in other cohorts. Due to the largely cross-sectional study design, we were unable to follow study volunteers beyond 6 months for IGRA-contacts, of whom only two converted to IGRA+ providing insufficient sample size to analyze correlates of IGRA conversion prospectively. Finally, all our biological samples in TB cohorts were derived from human blood and may not represent the abundance or function of NK cells within infected lung tissue. Future studies in *Mtb*-infected donors with multi-omic validation are needed to understand the role of CD8α^+^ NK cells at primary sites of infection.

In sum, our study adds to the growing literature highlighting a protective role for CD8α^+^ NK cells against both infections and non-infectious immune-mediated disease. Ongoing work will define how surface CD8αα^+^ NK cells may enhance targeting of *Mtb*-infected cells and will inform future cellular therapies and vaccination approaches to prevent or treat TB disease.

## MATERIALS AND METHODS

### Study design and participant recruitment

*Donor recruitment and protection of human subjects.* Haitian donors were enrolled through the Tri-Institutional Tuberculosis Research Unit (TBRU) at the Groupe Haitien d’etude du Sarcome de Kaposi et des Infections Opportunistes (GHESKIO) Centers under Institutional Review Board Approval at Weill Cornell Medicine and GHESKIO Centers in Haiti. All participants provided written informed consent, including consent for future use of biobanked samples for research. A dedicated clinical field team at the GHESKIO Centers in Port-au-Prince, Haiti, recruited as part of the NIH-funded TBRU (AI111143). LTBI was detected using QuantiFERON-TB® Gold (QIAGEN), and active TB was excluded by clinical screening for signs and symptoms of active TB including fever, cough, night sweats, weight loss, and abnormal findings on Chest X-ray. Most contacts were recruited close to the time of diagnosis of the index case and were living in the same house as the active TB case for at least 1 month in the 6 months prior to diagnosis (median time to contact recruitment from TB case diagnosis, 2.4 months; range, 0.2-26 months). To select for high transmission settings, each household was required to have a history of two active pulmonary TB cases. Healthy donors without reported TB exposure from the same community were recruited as controls. All cases with active pulmonary TB received periodic follow-up appointments while on treatment, and anyone with known contact with an active TB patient who remained IGRA-received 6-month follow-up and was rescreened with IGRA. The median smear grade of the active pulmonary TB cases in the households was 2-3+, conferring high risk for person-to-person transmission (*74*). All donors provided informed consent prior to peripheral blood donation for PBMC isolation. All donor samples were de-identified on site using a barcode system before they were shipped to Stony Brook University for analysis. Healthy donors were also recruited at Stony Brook University (SBU) under IRB2021-00478 (PI: Vorkas) and New York Blood Center (NYBC), which were used for stimulation assays and CITE-seq.

### PBMC isolation

PBMCs were isolated from peripheral blood at GHESKIO (Port-au-Prince, Haiti) using the Ficoll-Paque (GE Healthcare) density centrifugation, frozen in 5 × 10^6^ cells/ml aliquots in 90% FBS/10% DMSO (Thermo Fisher Scientific) and stored at –80°C prior to being shipped on dry ice to New York. PBMCs from healthy donors recruited at SBU and New York Blood Center (NYBC) were isolated using Ficoll-Paque, BD Vacutainer CPT cell preparation tube (Becton, Dickinson and Company) or SepMate tubes (Stem Cell Technologies) using the manufacturer protocol. PBMCs isolated at SBU were cryopreserved in serum free Bambanker medium (GC Lymphotec) and stored in liquid N_2_.

### Flow cytometry acquisition and analysis

A complete list of fluorescent antibodies is listed in Supplementary Table 1<u>: Reagents and resources</u>. All antibodies were commercially acquired, and multicolor panels were designed and validated using fluorescent minus one (FMO) controls. MR1 tetramers were obtained from the NIH Tetramer Core Facility (Atlanta, GA). In brief, cryopreserved PBMCs were thawed, plated, Fc receptor blocked (eBioscience), and underwent direct ex vivo extracellular staining for 15 minutes at room temperature (RT). For intracellular staining, PBMCs were incubated with 1X Brefeldin A (BioLegend) for final hour of incubation prior to staining with intracellular antibodies. Samples were then washed twice with FACS buffer and acquired using an Aurora Spectral Analyzer (Cytek). Flow cytometry analysis, including manual gating, was performed using FCS Express 7 (De Novo Software) or R (v4.3.1) with the following library packages downloaded from Bioconductor: *flowWorkspace, ggcyto, and flowCore*.

### *Mtb* whole cell lysate

*Mtb* whole cell lysate was prepared in Biosafety Level 3 facility at SBU. Fifty mL of *Mtb* Erdman 7H9 culture was grown to OD_600_ 0.6, centrifuged at 4000 × g for 10 min at RT and pellet was resuspended in 2.5 ml of 10mM Tris-HCl buffer (pH 7.5) and rotated for 2 hr at RT. The mixture was centrifuged at 500 × g for 15 min at RT and aqueous phase was collected and filtered twice through 0.22 µM spin filter (Corning Costar Spin-X) at 6000 × g for 5 min. The aliquots were stored at –80°C for further use.

### NK cell co-culture with K562 cell line

K562 cells (ATCC) were incubated in Iscove’s Modified Dulbecco’s Medium (IMDM) supplemented with 10% FBS. Cryopreserved K562 cells were thawed at 37°C water bath and cells were cultured at 1 x 10^5^ viable cells/ml in T-75 flask as per manufacturers protocol. Once the cell density reached 1 x 10^6^ cells/ml, K562 was co-incubated with PBMCs at 1:20 K562 to PBMC in a round bottom clear plate (Greiner) for 15 hrs, at 37°C with 5% CO_2_. At the beginning of incubation period anti-CD107a antibody was added to the culture. After incubation was complete, cells were washed, stained and acquired by flow cytometry.

### NK cell stimulation assay

Cryopreserved healthy donor PBMCs from SBU and NYBC were used for NK cell stimulation assays. Thawed PBMCs were cultured 1 x 10^4^ cells per well in round bottom clear plate (Greiner) for 15 hr, at 37°C with 5% CO_2_. Samples were stimulated and incubated overnight in complete RPMI media with 10% heat-inactivated FBS and supplemented with IL-2 (50 U/ml; PeproTech).

The cytokine stimulation conditions were performed with commercially available recombinant IL-12 (100 ng/mL; BioLegend), recombinant IL-18 (100 ng/mL; Biolegend) and IL15-Su construct (1 µg/ml) containing a truncated Sushi domain of IL15Rα synthesized by the Cheung lab, that is demonstrated to be a more potent and stable compound with longer half-life than recombinant IL15 (*52*). For *Mtb* stimulation condition we used 1:100 and 1:200 dilution of *Mtb* whole cell lysate.

### MHC I blockade

Cryopreserved healthy donor PBMCs from SBU and NYBC were used for MHC I blockade. PBMCs were thawed at 37°C and incubated in complete RPMI media with 10% heat-inactivated FBS and supplemented with IL-2 (50 U/ml; PeproTech). 1 x 10^4^ PBMCs were plated per well in round bottom clear plate (Greiner) and incubated with or without 5 µg/ml *Invivo*MAb anti-human MHC I (clone W6/32; Bio X cell) for 15 hr, at 37°C with 5% CO_2_ before being stained and acquired.

### scRNA-seq and CITE-seq library generation and sequencing

PBMCs were stained with fluorescent antibodies for cell sorting of MAIT cells (CD3^+^, CD161^+^, MR1-5-OP-RU tetramer^+^), NK cells (CD3^-^, CD19^-^, CD14^-^, CD16^+^ CD56^dim^), γδ T cells (CD3^+^, TCRγδ^+^) and iNKT cells (CD161^+^ iNKT^+^) using FACS. The sorted cells were blocked with human Trustain, FcX blocking reagent (BioLegend) for 10 minutes at 4°C and then stained with TotalSeq-C barcoded tagged and hashtags antibodies for 30 minutes at 4°C in a total volume of 100ul. The residual TotalSeq-C and hashtags antibodies were removed by centrifugation at 400 *g* for 5 minutes at 4°C. A complete hashtag and TotalSeq-C antibody list is provided in **Supplementary Table 1: Reagent and Resources.**

Gene expression and surface antigen library preparation was conducted at the Stony Brook University Single Cell Genomics Facility. Briefly, cell viability and concentrations were determined on a Countess 3FL with trypan blue staining. Approximately 5,000 live cells from each condition were loaded in a single 10X Genomics 5’ NextGEM Single Cell assay reaction (30,000 total cells per reaction), to perform 5’ single cell immune profiling assay using a Chromium iX instrument (10X Genomics). Gene expression (GEX) and surface antigen (SA) libraries were prepared, and quality controlled using Chromium NextGEM 5’ Single Cell with Feature Barcode assay and a TapeStation 4200, according to the manufacturer’s instructions. The libraries were sequenced at a depth of approx. 25,000 (GEX) and 10,000 (SA) paired reads/cell on an Illumina NovaSeqX sequencer (Novogene, Inc).

The gene expression and surface feature barcode raw FASTQ files of single cell RNA-seq were analyzed on Cell Ranger (10x Genomics, v.7.0) (https://www.10xgenomics.com/analysis-guides/demultiplexing-and-analyzing-5%E2%80%99-immune-profiling-libraries-pooled-with-hashtags), multi-pipeline with human reference transcriptome (GRCh38) downloaded from 10x Genomics website and hashtag configuration file, which contained the barcodes related to each different samples.

### Single-cell RNAseq analysis

The generated gene expression matrix files were analyzed with R (4.3.2) using the *Seurat* (5.0) package and cells were filtered based on following criteria: 1.) the number of genes was equal to or greater than 200 and below than 6000 and 2.) the mitochondrial percentage was less than 5. Using the FindVariableFeatures() function we selected the top 3000 features with highest variance. Then, different samples were integrated using the harmony algorithm and scaled. Log-normalization and principal component analysis (PCA) was done on integrated values. The top 20 principal components were selected for clustering and visualization with UMAP using the default Louvain algorithm. Differential gene expression was performed using FindAllMarkers() with a minimum threshold of 25% cells expressing.

### Statistical Analysis

Statistical analysis was performed using R (4.3.2) and GraphPad Prism (10.1.2). Non-parametric testing with Wilcoxon tests was used to compare the means between groups for flow markers. P-values were not adjusted because cross-wise comparisons were not performed between all six clinical groups. Only comparisons a priori hypothesized to represent biological features were tested, these included: IGRA-contacts vs controls, IGRA+ contacts vs controls, IGRA-contact vs IGRA+ control, and IGRA+ control vs Active TB.

### Supplementary Materials

Supplementary Table 1: Reagents and resources. See attached file.

## Acknowledgments

We are thankful for the support of the Stony Brook Foundation, the Department of Medicine of the Renaissance School of Medicine and the Office of the Vice President of Research of Stony Brook University. We thank Daniel W. Fitzgerald, MD (Weill Cornell Medicine, Center for Global Health) and Michael S. Glickman (Memorial Sloan Kettering Cancer Center) for generously providing clinical samples from TB contact cohorts. We thank Joseph C. Sun (MSKCC) for valuable discussions on adaptive NK cells. We are also grateful to Thomas MacCarthy, PhD (Stony Brook University, Applied Mathematics) for supportive advice regarding CITE-seq analyses, who passed away during the preparation of this manuscript.

## Funding

This study was supported by the Tri-I TBRU, part of the TBRU Network (U19 AI11143). CKV acknowledges support from the NIAID T32 Pathogenesis of Infectious Diseases Training Program (T32AI007613-18; Roy Gulick, WCM), the IDSA Education and Research Foundation Postdoctoral Fellowship Award in Infectious Diseases, a Potts Memorial Foundation Award, NIAID K08AI132739, R21AI171578, and R21AI183259.

## Author contributions

The following authors contributed to the manuscript in various ways. Each author’s contribution is listed below.

Conceptualization: KH, JBL, CKV

Methodology: NKC, AP, RK, KCH, JBL, CKV

Investigation: NM, AP, RK, TK, BR, NKC, HX, JBL and CKV

Visualization: NM, AP, RK, JB, BR, DEC, JBL, and CKV

Funding acquisition: CKV

Writing – original draft: NM, AP, CKV

Writing – review & editing: all authors

## Competing interests

Authors declare that they have no competing interests.

## Data and materials availability

All data, code, and materials used in the analysis will be available to any researcher for purposes of reproducing or extending the analysis.

## Notes

### Competing Interest Statement

The authors have declared no competing interest.

### Summary of Updates

The author affiliations for have been updated.

