## Supplemental Figures for "Loss of circulating CD8α^+^ NK cells during human *Mycobacterium tuberculosis* infection"

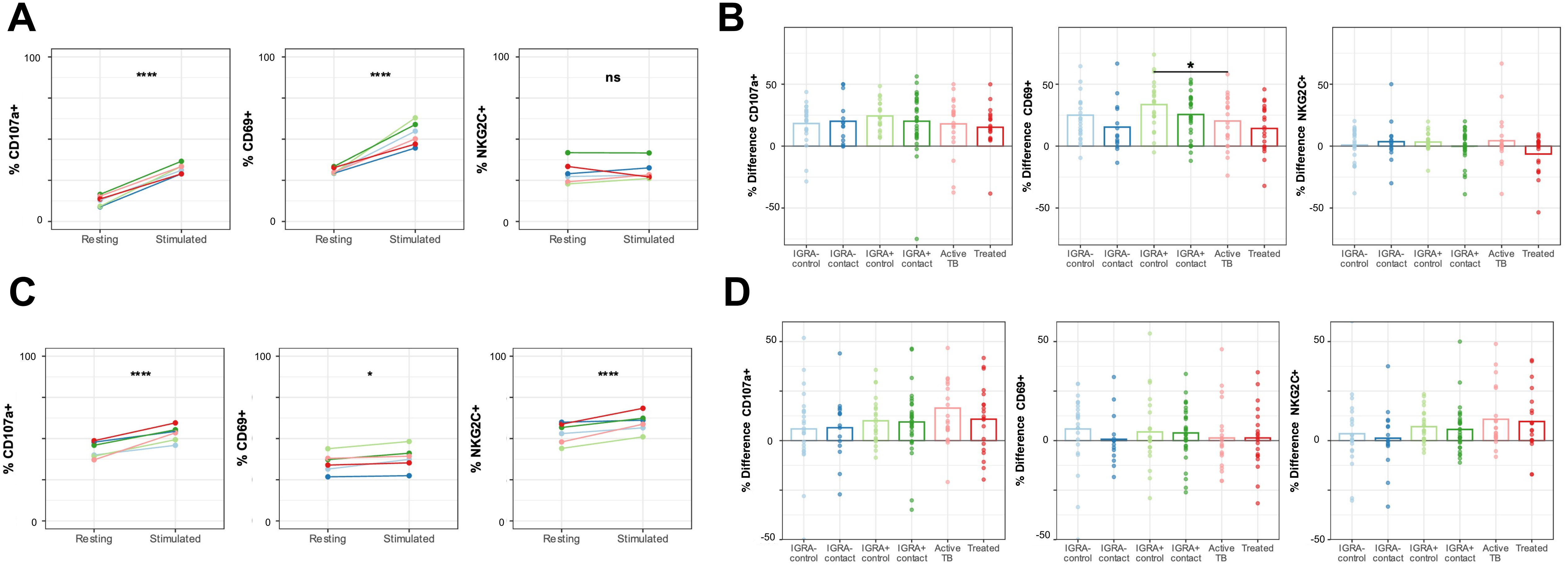


**Fig. S1. CD56$^{bright}$ and CD56$^{-}$ NK cell response to K562 re-stimulation measured by CD69, CD107a, and NKG2C surface expression. (A)** Overall results of activation assay for CD56$^{bright}$ NK cells for each activation marker. Y-axis represents mean percent positive. P-values calculated as a global paired Wilcoxon test including all clinical groups. Each clinical group is displayed separately indicated by line color. **(B)** Percent difference (stimulated minus resting) of three activation markers in CD56$^{bright}$ NK cells comparing between clinical group. Brackets represent Wilcoxon tests with unadjusted p-values. **(C)** Overall results of activation assay for CD56$^{-}$ NK cells, as in panel A. **(D)** Percent difference (stimulated minus resting) of three activation markers in CD56$^{-}$ NK cells comparing between clinical group. Brackets represent Wilcoxon tests with unadjusted p-values. $^{*}p \leq 0.05$, $^{****}p \leq 0.0001$.

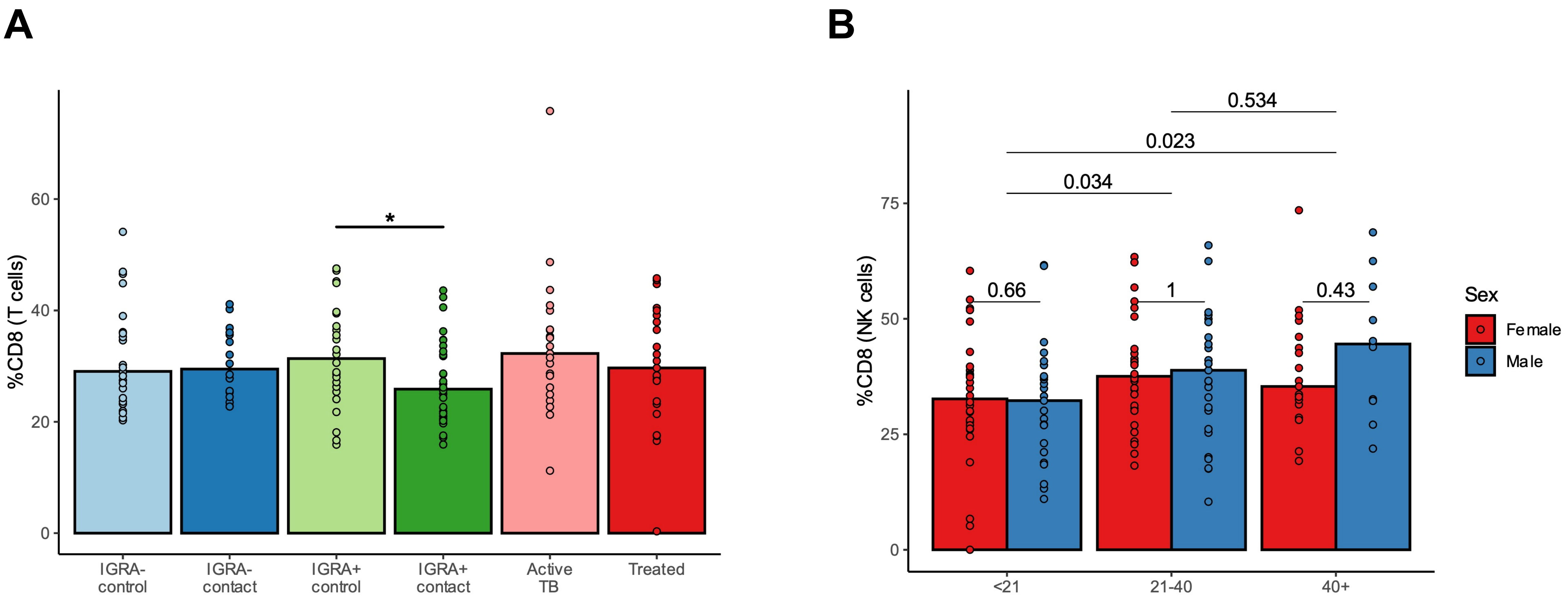


**Fig. S2. CD8α expression in T cells. (A)** Bar graph demonstrating frequency of T cells that are CD8α positive compared between clinical groups. Brackets represent Wilcoxon tests. **(B)** Bar graph demonstrating frequency of CD8α$^+$ NK cells among age and sex demographics. Color represents sex, with three age ranges displayed. Brackets represent Wilcoxon tests. *p ≤ 0.05.

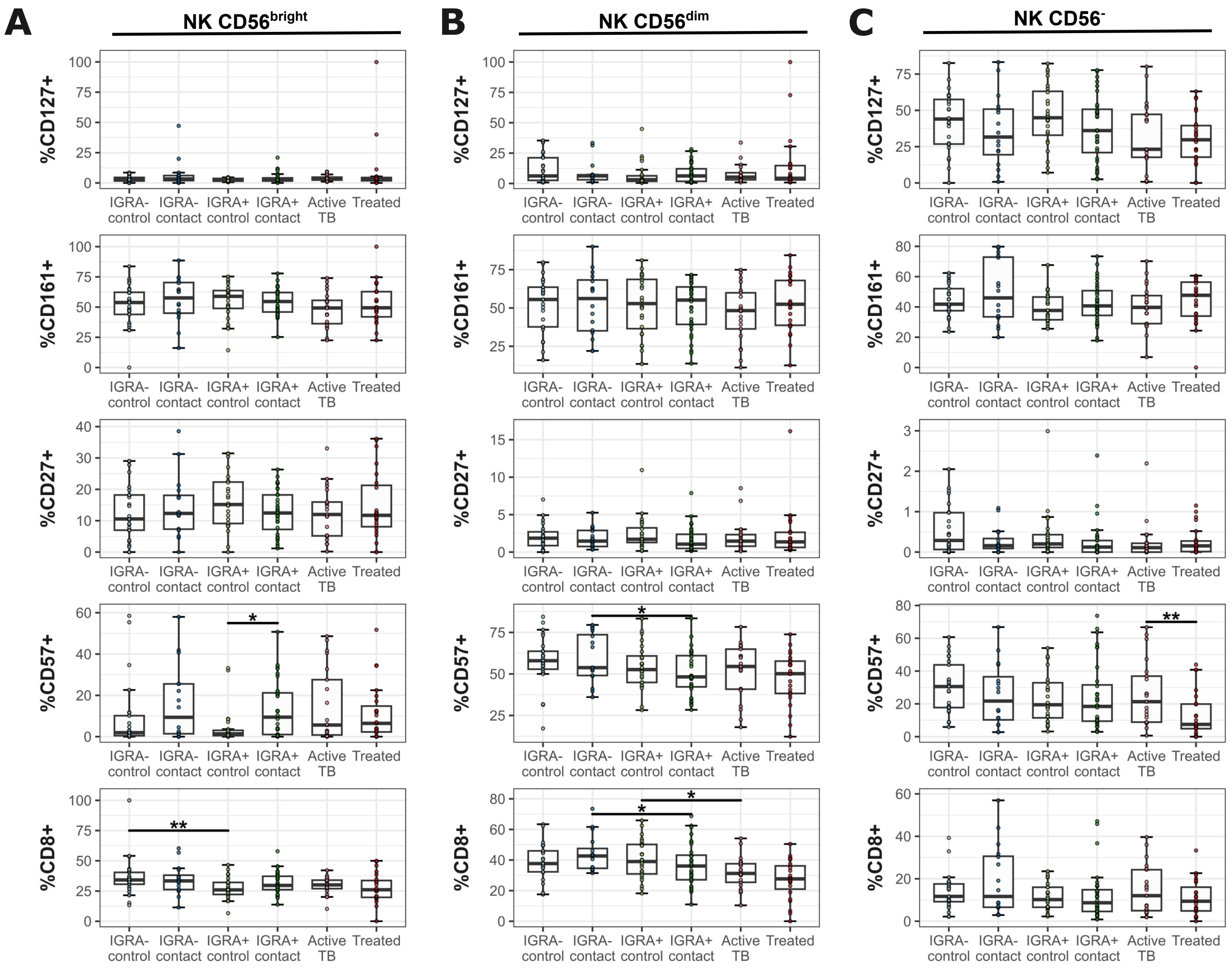


**Fig. S3. Baseline surface expression for potential memory markers among NK cell subsets in ex vivo stained samples. (A)** Percent positive of five putative memory markers in CD56$^{bright}$ NK cells. Brackets represent Wilcoxon tests with unadjusted p-values. **(B)** Percent positive of five putative memory markers in CD56$^{dim}$ NK cells. Brackets represent Wilcoxon tests with unadjusted p-values. **(C)** Percent positive of five putative memory markers in CD56$^{-}$ NK cells. Brackets represent Wilcoxon tests with unadjusted p-values. *$p \leq 0.05$, **$p \leq 0.01$.

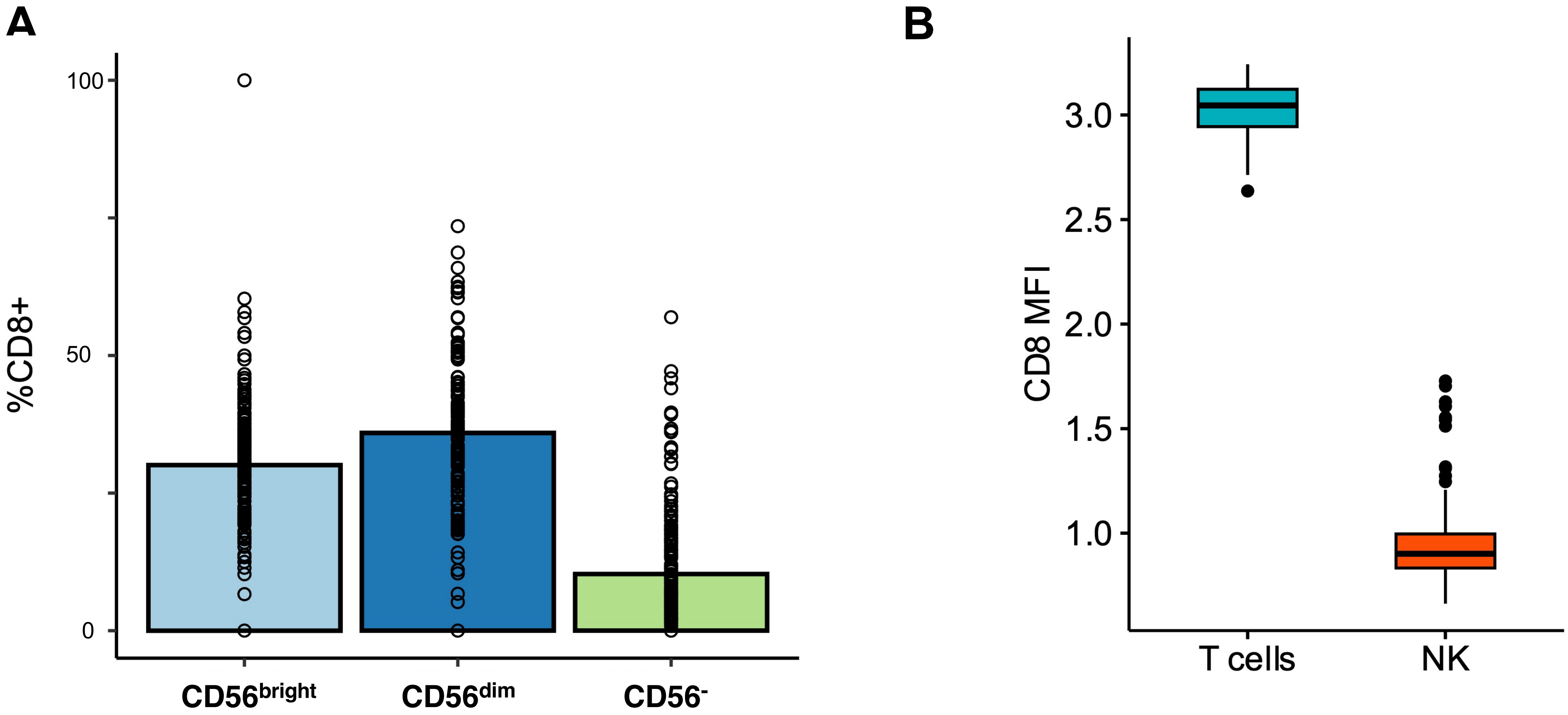


**Fig. S4. Expression of CD8α among three NK cell subsets: CD56$^{bright}$, CD56$^{dim}$, and CD56$^{-}$ (A)** Median CD8α frequency across CD56 stratified NK cell subsets (**B)** Boxplot of MFI values between CD8α$^{+}$ NK cells versus CD8α$^{+}$ T cells.

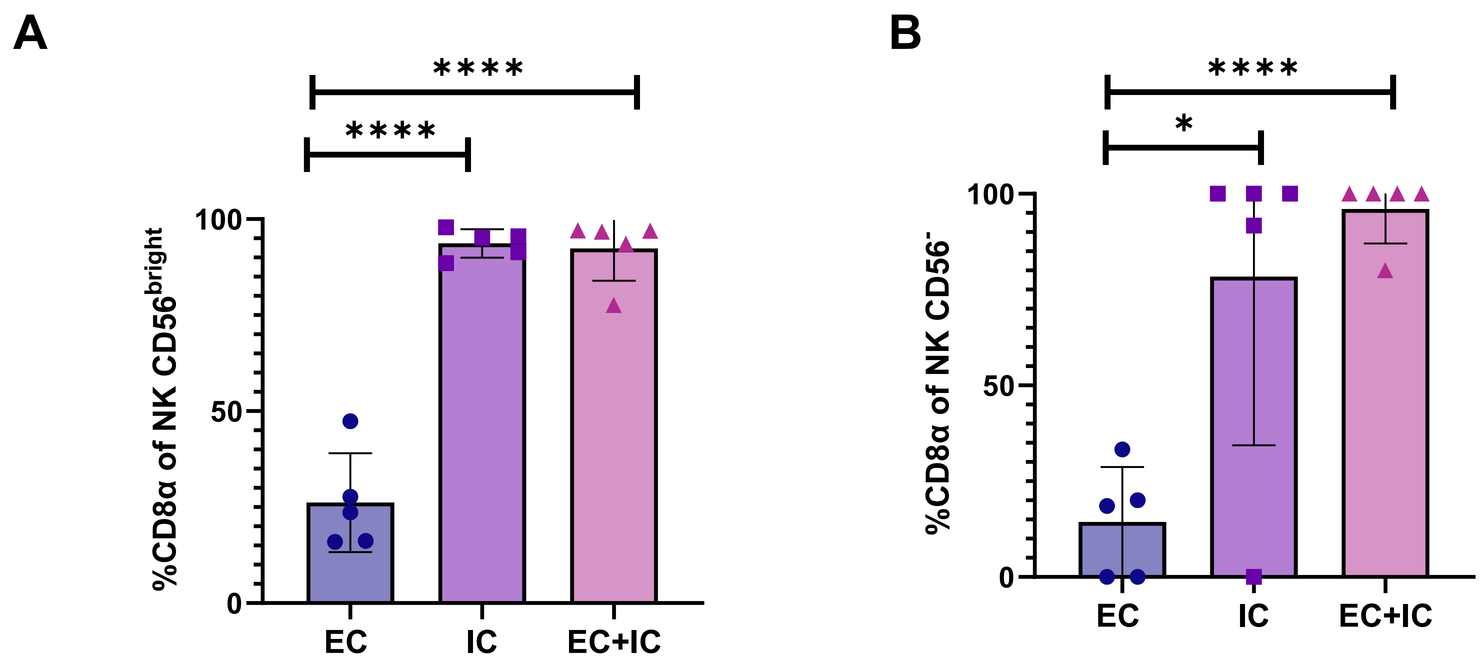


**Fig. S5. Ubiquitous intracellular CD8α expression among CD56⁻ and CD56$^{bright}$ NK cells. (A)** Percentage of CD8α$^+$ cells in CD56$^{bright}$ NK cells. Brackets represent unpaired t-tests. **(B)** Percentage of CD8α$^+$ cells in CD56$^-$ NK cells. Brackets represent unpaired t-tests. EC (**e**xtra**c**ellular/surface), IC (**i**ntra**c**ellular), and EC + IC (both extracellular and Intracellular). *p<0.05, ****p<0.0001.

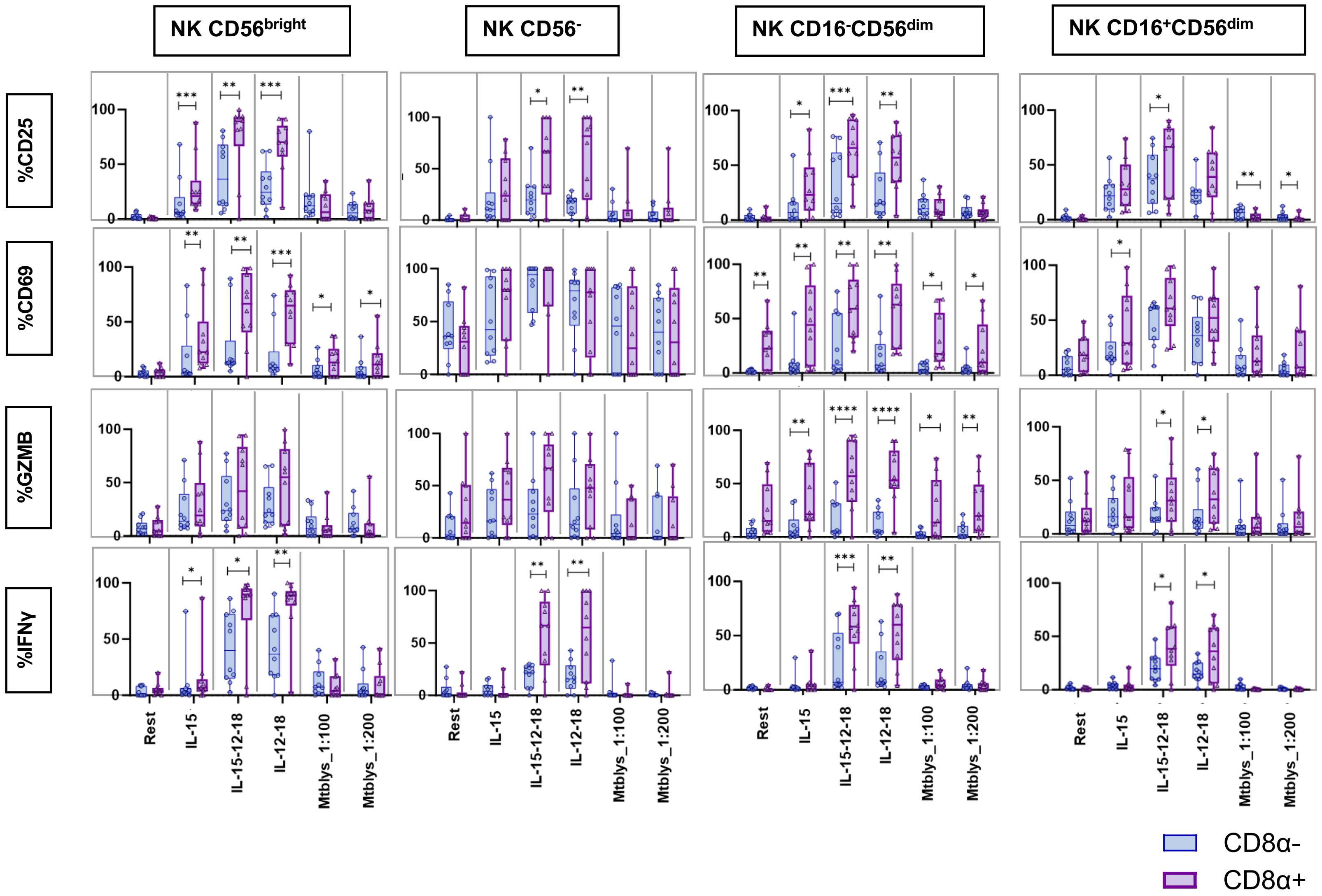


**Fig. S6. Enhanced responsiveness of CD8α$^{+}$ NK subsets upon cytokine stimulation. (A)** NK cells were cultured for 15 hours in either plain media or supplemented with IL-15, IL15/12/18, IL-12/18, or MtbLys at dilutions of 1:100 and 1:200. Percent positive of CD25, CD69, granzyme B (GzmB) and interferon gamma (IFNγ) in CD8α$^{+}$ and CD8α$^{-}$ CD56$^{bright}$, CD56$^{-}$, CD16$^{+}$ and CD16$^{-}$ NK CD56$^{dim}$ populations measured by flow cytometry. *$p \leq 0.05$, **$p \leq 0.01$, ***$p \leq 0.001$ ****$p \leq 0.0001$

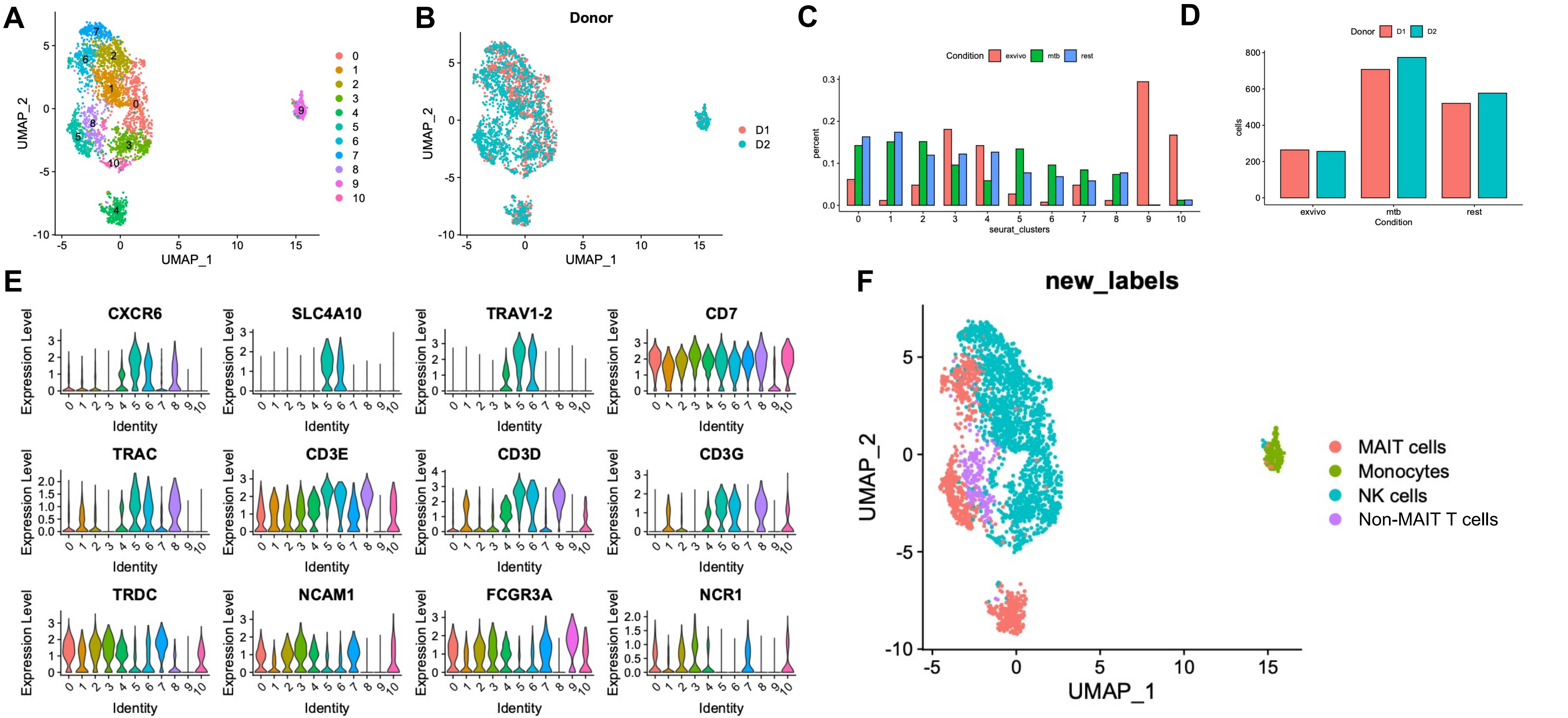


**Fig. S7. CITE-seq clustering of sorted MAIT cells, iNKT, NK cells, and γδ T cells in three stimulation conditions**. **(A)** UMAP representation of Louvain clustering output. **(B)** Cells labeled by donor origin. **(C)** Bar graph of total number of cells per cluster separated by condition. **(D)** Total cells per donor and condition. **(E)** Expression of select markers comparing original cluster identities. **(F)** Final annotation of clusters after merging of similar clusters.

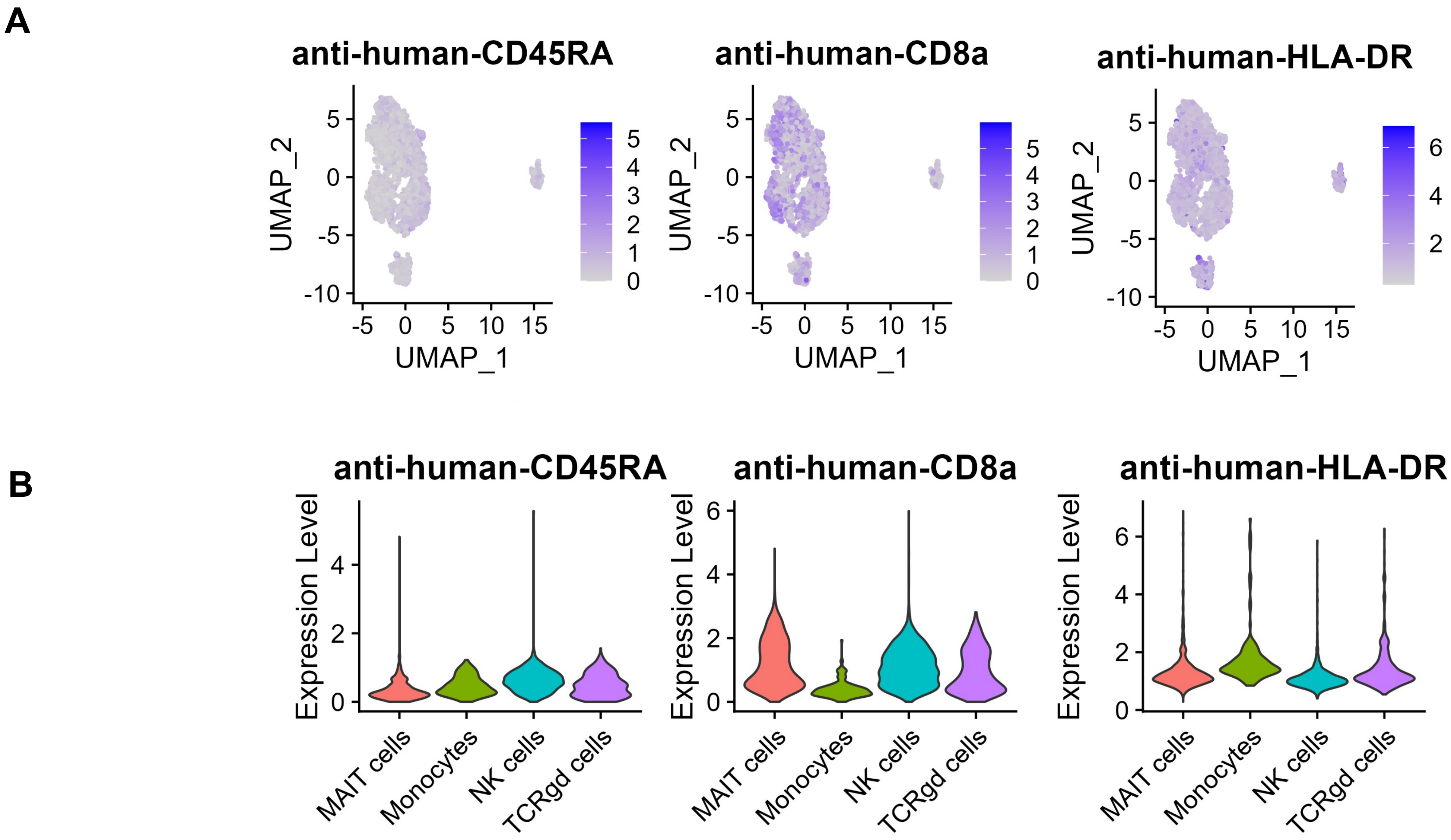


**Fig. S8. CITE-seq of innate lymphocytes analyzed by surface barcode antibody detection in annotated clusters of MAIT cells, NKT, NK cells, and γδ T cells. (A)** UMAP representation of all cells colored by surface expression of marker stated in the plot title. **(B)** Violin plot of the expression level of four selected markers between each cell subset after manual annotation.
