## Supplemental Table 1 for "Loss of circulating CD8α^+^ NK cells during human *Mycobacterium tuberculosis* infection"

**Supplementary Table 1: Reagents and Resources**

| REAGENT or RESOURCE | SOURCE | IDENTIFIER |
|---|---|---|
| **Antibodies (anti-human)** | | |
| BV421 CD69 (clone: FN50) | BioLegend | Cat#310930; RRID:AB_2561909 |
| BV570 CD16 (clone: 3G8) | BioLegend | Cat#302036; RRID:AB_2632790 |
| BV605 TCRVα7.2 (clone:3C10) | BioLegend | Cat#351720; RRID:AB_2563991 |
| BV711 CD25 (clone: BC96) | BioLegend | Cat#302636; RRID:AB_2562910 |
| BV750 CD56 (clone: B159) | BD | Cat#747179; RRID: AB_2871913 |
| BV785 IFN-γ (clone: 4S.B3) | BioLegend | Cat#502542; RRID: AB_2563882 |
| FITC Granzyme B (clone: GB11) | BioLegend | Cat#515403; RRID:AB_2114575 |
| cFluor YG584 CD4 (clone: SK3) | Cytek | Cat#R7-20041 |
| PerCP CD4 (clone: SK3) | BioLegend | Cat#980820 |
| PE MR1 (clone: 26.5) | BioLegend | Cat#361106; RRID: AB_2563043 |
| APC CD161 (clone: DX12) | BD | Cat#550968; RRID:AB_398482 |
| AlexaFluor700 CD3 (clone: UCHT1) | BD | Cat# 557943; RRID:AB_396952 |
| APC-Cy7 CD8 (clone: SK1) | BD | Cat#560179; RRID:AB_1645481 |
| PE-Cy7 CD8b (clone: SIDI8BEE) | Invitrogen | Cat#25527341; RRID: AB_11217487 |
| SparkNIR685 CD19 (clone:HIB19) | BioLegend | Cat#302270; RRID:AB_2832581 |
| PE-Vio770 CD158i_KIR2DS4 (clone:REA860) | Miltenyi Biotec | Cat#130114617; RRID:AB_2655372 |
| Fc Receptor Binding Inhibitor Polyclonal Antibody | eBioscience | Cat#14-9161-73; RRID:AB_468582 |
| Zombie red Fixable viability kit | BioLegend | Cat# 423110 |
| **Sorting fluorophore antibodies** | | |
| APC CD161 (clone: DX12) | BD Pharmingen | Cat# 550968; RRID:AB_398482 |
| PE MR1 tetramer | PE | |
| Alexa Fluor 700 CD3 (clone: UCHT1) | BD Biosciences | Cat# 557943; RRID:AB_396952 |
| BV480 CD14 (clone: M5E2) | BD Pharmingen | Cat# 746304; RRID:AB_2743629 |
| PE-Cy5 CD19 (clone: HIB19) | Biolegend | Cat# 302210; RRID: AB_314240 |
| FITC CD16 (clone: B73.1) | Biolegend | Cat# 360716; RRID: AB_2563071 |
| PE-Cy7 CD56 (clone: CMSSB) | Fisher | Cat# 25-0567-42; RRID:AB_11041529 |
| BV650 TCRgd (clone: B1.1) | BD Pharmingen | Cat# 564156; RRID:AB_2738628 |
| BV510 TCRVa24 (iNKT) (clone: 6B11) | Biolegend | Cat# 342918; RRID: AB_2564006 |
| **Hashtag antibodies** | | |
| **Antibody (Sequence) (Clone)** | | |
| TotalSeq-C0251 MB_Exvivo (GTCAACTCTTTAGCG) (LNH-94;2M2) | BioLegend | Cat# 394661; RRID:AB_2801031 |
| TotalSeq-C0252 MB_Rest (TGATGGCCTATTGGG) (LNH-94;2M2) | BioLegend | Cat# 394663; RRID:AB_2801032 |
| TotalSeq-C0253 MB_Mtblysate (TTCCGCCTCTCTTTG) (LNH-94;2M2) | BioLegend | Cat# 394665; RRID:AB_2801033 |
| TotalSeq-C0254 HD_073409_Exvivo (AGTAAGTTCAGCGTA) (LNH-94;2M2) | BioLegend | Cat# 394667; RRID:AB_2801034 |
| TotalSeq-C0255 HD_073409_Rest (AAGTATCGTTTCGCA) (LNH-94;2M2) | BioLegend | Cat# 394669; RRID:AB_2801035 |
| TotalSeq-C0256 HD_073409_Mtblysate (GGTTGCCAGATGTCA) (LNH-94;2M2) | BioLegend | Cat# 394671; RRID:AB_2801036 |
| **TotalseqC antibodies** | | |
| TotalSeq-C0072 CD4 (TGTTCCCGCTCAACT) (RPA-T4) | BioLegend | Cat# 300567; RRID:AB_2800725 |
| TotalSeq-C0390 CD127 (GTGTGTTGTCCTATG) (A019D5) | BioLegend | Cat# 351356; RRID:AB_2800937 |
| TotalSeq-C0081 CD14 (TCTCAGACCTCCGTA) (M5E2) | BioLegend | Cat# 301859; RRID:AB_2800736 |
| TotalSeq-C0396 CD26 (GGTGGCTAGATAATG) (BA5b) | BioLegend | Cat# 302722; RRID:AB_2810435 |
| TotalSeq1-C0084 CD 56 (NCAM) (TTCGCCGCATTGAGT) (QA17A16) | BioLegend | Cat# 392425; RRID:AB_2801024 |
| TotalSeq1-C0050 CD19 (CTGGGCAATTACTCG) (HIB19) | BioLegend | Cat# 302265; RRID:AB_2800741 |
| TotalSeq-C0147 CD62L (GTCCCTGCAACTTGA) (DREG-56) | BioLegend | Cat# 304851; RRID:AB_2800770 |
| TotalSeq-C0155 CD107a (LAMP-1) (CAGCCCACTGCAATA) (H4A3) | BioLegend | Cat# 328649; RRID:AB_2800854 |
| TotalSeq-C0149 CD161 (GTACGCAGTCCTTCT) (HP-3G10) | BioLegend | Cat# 339947; RRID:AB_2810532 |
| TotalSeq-C0158 CD134 (OX40) (AACCCACCGTTGTTA) (Ber-ACT35 (ACT35) | BioLegend | Cat# 350035; RRID:AB_2800932 |
| TotalSeq-C0101 CD335 (NKg46) (ACAATTTGAACAGCG) (9 E2) | BioLegend | Cat# 331941; RRID:AB_2800874 |
| TotalSeq-C0171 CD278 (ICOS) (CGCGCACCCATTAAA) (C398.4A) | BioLegend | Cat# 313553; RRID:AB_2800874 |
| TotalSeq-C0034 CD3 (CTCATTGTAACTCCT) (UCHT1) | BioLegend | Cat# 300479; RRID:AB_2800823 |
| TotalSeq-C0146 CD69 (GTCTCTTGGCTTAAA) (FN50) | BioLegend | Cat# 310951; RRID:AB_2800810 |
| TotalSeq-C0046 CD8 (GCGCAACTTGATGAT) (SK1) | BioLegend | Cat# 344753; RRID:AB_2800922 |
| TotalSeq-C0159 HLA-DR (AATAGCGAGCAAGTA) (L243) | BioLegend | Cat# 307663; RRID:AB_2800795 |
| TotalSeq-C0153 KLRG-1 (MAFA) (CTTATTTCCTGCCCT) (SA231A2) | BioLegend | Cat# 367737; RRID:AB_2904401 |
| TotalSeq-C0063 CD45RA (TCAATCCTTCCGCTT) (HI100) | BioLegend | Cat# 304163; RRID:AB_2800764 |
| TotalSeq-C0080 CD8a (GCTGCGCTTTCCATT) (RPA-T8) | BioLegend | Cat# 301071; RRID:AB_2800730 |
| TotalSeq-C0007 CD274 (B7-H1, PD-L1) (GTTGTCCGACAATAC) (29E.2A3) | BioLegend | Cat# 329751; RRID:AB_2800860 |
| TotalSeq-C0032 CD154 (GCTAGATAGATGCAA) (24-31) | BioLegend | Cat# 310849; RRID:AB_2800808 |
| TotalSeq-C0053 CD11c (TACGCCTATAACTTG) (S-HCL-3) | BioLegend | Cat# 371521; RRID:AB_2801018 |
| TotalSeq-C0083 CD16 (AAGTTCACTCTTTGC) (3G8) | BioLegend | Cat# 302065; RRID:AB_2800738 |
| TotalSeq-C0867 CD94 (CTTTCCGGTCCTACA) (DX22) | BioLegend | Cat# 305523; RRID:AB_2814143 |
| TotalSeq-C0420 CD158 (KIR2DL1/S1/S3/S5) (TATCAACCAACGCTT) (HP-MA4) | BioLegend | Cat# 339517; RRID:AB_2814252 |
| TotalSeq-C0592 CD158b (KIR2DL2/L3, NKAT2 (GACCCGTAGTTTGAT) (DX27) | BioLegend | Cat# 312619; RRID:AB_2819944 |
| TotalSeq-C0156 CD96 (Fas) (CCAGCTCATTAGAGC) (DX2) | BioLegend | Cat# 305651; RRID:AB_2800787 |
| TotalSeq-C0047 CD56 (NCAM) (TCCTTTCCTGATAGG) (5.1H11) | BioLegend | Cat# 362559; RRID:AB_2801002 |
| TotalSeq-C0151 CD152 (CTLA-4) (ATGGTTCACGTAATC) (BNI3) | BioLegend | Cat# 369621; RRID:AB_2801015 |
| TotalSeq-C0165 CD314 (NK2D) (CGTGTTTGTTCCTCA) (1D11) | BioLegend | Cat# 320837; RRID:AB_2800844 |

| | | |
|---|---|---|
| TotalSeq-C0145 CD103 (Integrin aE) (GACCTCATTGTGAAT) (Ber-ACT8) | BioLegend | Cat# 350233; RRID:AB_2800933 |
| TotalSeq-C0161 CD11b (GACAAGTGATCTGCA) (ICRF44) | BioLegend | Cat# 301359; RRID:AB_2800732 |
| TotalSeq-C0168 CD57 (AACTCCCTATGGAGG) (QA17A04) | BioLegend | Cat# 393321; RRID:AB_2801030 |
| TotalSeq-C0154 CD27 (GCACTCCTGCATGTA) (O323) | BioLegend | Cat# 302853; RRID:AB_2800747 |
| TotalSeq-C0599 CD158e1 (GGACGCTTTCCTTGA) (DX9) | BioLegend | Cat# 312725; RRID:AB_2814161 |
| TotalSeq-C0087 CD45RO (CTCCGAATCATGTTG) (UCHL1) | BioLegend | Cat# 304259; RRID:AB_2800766 |
| TotalSeq-C0391 CD45 (TTTGTCCTGTACGCC) (HI30) | BioLegend | Cat# 304068; RRID:AB_2800762 |
| TotalSeq-C0085 CD25 (TGCAATTACCCGGAT) (BC96) | BioLegend | Cat# 302649; RRID:AB_2800745 |
| **Fc Block** | | |
| Human Trustain FcX | BioLegend | Cat#422301; RRID: AB_2818986 |
| **Chemicals, peptides, and recombinant proteins** | | |
| Cytiva Ficoll-Paque™ PREMIUM | Cytiva | Cat#45001751 |
| Ficoll Paque Plus | GE Healthcare | Cat#17144002 |
| Fetal Bovine Serum (FBS) | Gibco | Cat#10437028 |
| Bambanker Serum-free cell freezing media | Lymphotec Inc. | Cat#9582225 |
| Brefeldin A solution (1000x) | Biolegend | Cat#420601 |
| Difco Middlebrook 7H9 broth | BD | Cat#271310 |
| Difco Middlebrook 7H10 Agar | BD | Cat#262710 |
| BD BBL Middlebrook OADC | BD | Cat#B12351 |
| Recombinant human IL12 | BioLegend | Cat#573002 |
| Recombinant human IL18 | BioLegend | Cat#592104 |
| Recombinant human IL2 | PeproTech | Cat#200-02 |
| Recombinant human IL2 | Gibco | Cat#PHC0021 |
| Recombinant human IL2 (hIL-2) | Roche | Cat#HIL2-RO |
| Penicillin/Streptomycin | Gibco | Cat#15-140-122 |
| IMDM | Gibco | Cat#12440053 |
| RPMI 1640 | Gibco | Cat#21870092 |
| L-glutamine | Gibco | Cat#25030149 |
| Fetal Bovine Serum | Gibco | Cat#26140079 |
| HEPES | Gibco | Cat#15630080 |
| Sodium pyruvate | Gibco | Cat#11360070 |
| MEM Nonessential amino acids | Gibco | Cat#11140050 |
| Flow Cytometry Staining Buffer | eBioscience | Cat#00-4222-26 |
| Fixation/Permeabilization concentrate | eBioscience | Cat#00-5123-43 |
| Permeabilization Buffer | eBioscience | Cat#00-8333-56 |
| 2-mercaptoethanol | Sigma | Cat#M6250-250ML |
| **Biological samples** | | |
| Healthy human PBMCs | NYBC, SBU | N/A |
| Healthy human TB contact/control PBMCs | GHESKIO Centers | N/A |
| **Software and algorithms** | | |
| Cellranger | v.7.0 | https://www.10xgenomics.com/support/software/cell-ranger/latest/analysis/running-pipelines/cr-3p-multi |
| Seurat | v.5.0.1 | https://satijalab.org/seurat/articles/install.html |
| R | v.4.3.1 | https://www.r-project.org/ |
| FCS express v7 | DeNovo Software | https://denovosoftware.com/ |
| Prism v11 | Graphpad Software | https://www.graphpad.com/scientific-software/prism/ |
